# New SHIVs and Improved Design Strategy for Modeling HIV-1 Transmission, Immunopathogenesis, Prevention and Cure

**DOI:** 10.1101/2021.01.13.426578

**Authors:** Hui Li, Shuyi Wang, Fang-Hua Lee, Ryan S. Roark, Alex I. Murphy, Jessica Smith, Chengyan Zhao, Juliette Rando, Neha Chohan, Yu Ding, Eunlim Kim, Emily Lindemuth, Katharine J. Bar, Ivona Pandrea, Christian Apetrei, Brandon F. Keele, Jeffrey D. Lifson, Mark G. Lewis, Thomas N. Denny, Barton F. Haynes, Beatrice H. Hahn, George M. Shaw

## Abstract

Simian-human immunodeficiency virus (SHIV) chimeras contain the HIV-1 envelope (*env*) gene embedded within an SIVmac proviral backbone. Previously, we showed that substitution of Env residue 375-Ser by bulky aromatic residues enhances Env binding to rhesus CD4 and enables primary or transmitted/founder (T/F) HIV-1 Envs to support efficient SHIV replication in rhesus macaques (RMs). Here, we test this design strategy more broadly by constructing and analyzing SHIVs containing ten strategically selected primary or T/F HIV-1 Envs corresponding to subtypes A, B, C, AE and AG, each with six allelic variants at position 375. All ten SHIVs bearing wildtype Env375 residues replicated efficiently in human CD4^+^ T cells, but only one of these replicated efficiently in rhesus CD4^+^ T cells. This was a SHIV whose subtype AE Env naturally contained a bulky aromatic His residue at position 375. Replacement of wildtype Env375 residues by Trp, Tyr, Phe or His in the other nine SHIVs uniformly led to efficient replication in rhesus CD4+ T *in vitro* and in RMs *in vivo*. Env375-Trp – the residue found most frequently among SIV strains infecting Old World monkeys – was favored for SHIV replication in RMs, although some SHIVs preferred Env375-Tyr, -His or -Phe. Nine SHIVs containing optimized Env375 alleles were grown large scale in primary activated rhesus CD4^+^ T cells to serve as challenge stocks in preclinical prevention trials. These virus stocks were genetically homogeneous, native-like in Env antigenicity and tier-2 neutralization sensitivity, transmissible by rectal, vaginal, penile, oral or intravenous inoculation routes, and exhibited acute and early replication kinetics that were indistinguishable from HIV-1 infection in humans. Finally, to expedite future SHIV constructions and eliminate short redundant elements in *tat1* and *env* gp41 that were spontaneously deleted in chronically infected monkeys, we engineered a simplified second-generation SHIV design scheme and validated it in RMs. Overall, our findings demonstrate that SHIVs bearing primary or T/F Envs with bulky aromatic amino acid substitutions at position Env375 consistently replicate in RMs, recapitulating many features of HIV-1 infection in humans. We further show that SHIV challenge stocks grown in primary rhesus CD4^+^ T cells are efficiently transmitted by mucosal routes common to HIV-1 infection and can be used effectively to test for vaccine efficacy in preclinical monkey trials.

## Introduction

Simian-human immunodeficiency virus (SHIV) infection of Indian rhesus macaques (RMs) is an important outbred animal model for studying HIV-1 transmission, prevention, immunopathogenesis and cure (1–3). Such research is especially timely, given recent progress with active and passive immunization (4–11) and novel approaches to HIV-1 cure (https://www.niaid.nih.gov/diseases-conditions/hiv-cure-research) (12–18), all of which can benefit from rigorous testing and iterative refinement in animal models. Given the multifaceted roles of HIV-1 envelope (Env) in cell tropism and virus entry, and as a target for neutralizing and non-neutralizing antibodies, the particular features of HIV-1 Envs that are selected for SHIV construction and analysis are of paramount importance. This is especially true for vaccine studies designed to administer (10, 11) or elicit (6, 19, 20) broadly neutralizing antibodies (bNAbs).

SHIVs have a long history dating to 1992 when Sodroski and colleagues first subcloned the *tat*, *rev* and *env* sequences HIV-1 HXB2c into SIVmac239 (21). This clone was further modified by substitution of the *env* from the dual CCR5/CXCR4 tropic HIV-1 89.6 strain and later adapted by serial passage in RMs, eventually yielding the molecular clone SHIV-KB9 (22). Thus, the earliest SHIVs contained T-cell line adapted, *in vivo* passaged HIV-1 Envs that were CXCR4 tropic, highly syncytium-inducing and cytopathic, and led to accelerated disease in monkeys. As a consequence, many of the essential features of HIV-1 biology, including cell and tissue tropism, sensitivity to neutralizing antibodies (NAbs), immunopathogenesis, transmission efficiency and natural history, were not faithfully represented in the macaque model (3). Attempts to develop a SHIV infection model that included primary (non-T-cell line adapted) CCR5-tropic Envs were generally met with failure, and when they were successful, such SHIVs often required adaptation by serial monkey passage to achieve consistent replication *in vivo* (3, 23–25). In an attempt to better understand restrictions to SHIV infection and replication in RMs, Overbaugh and Sawyer examined the affinity of primary HIV-1 Envs to rhesus CD4 (26, 27). They discovered that the Envs of most primary HIV-1 strains exhibited low affinity for rhesus CD4 and did not support efficient virus entry into rhesus cells. Overbaugh identified a key amino acid at position 39 in domain 1 of rhesus CD4 that differed between human and rhesus CD4 and was largely responsible for the poor binding and infectivity of primary HIV-1 Envs in rhesus cells (27). This presented a major obstacle to new SHIV designs. Hatziioannou identified a mutation at residue 281 in the CD4 binding region of HIV-1 Env that occurred commonly in SHIV-infected RMs, where it could be shown to facilitate virus replication (28). However, unlike the Env375 substitution, the 281 substitution on its own was unable to consistently convert primary or transmitted/founder (T/F) Envs, which fail to replicate efficiently in RMs, to do so. Moreover, the addition of the 281 mutation to SHIV Envs that already contain a rhesus-preferred Env375 allele, did nothing to further enhance virus replication in rhesus animals (29).

We noted from studies by Finzi and Sodroski (30) that residue 375 in the CD4 binding pocket of primate lentiviral Envs was under strong positive evolutionary pressure across the broad spectrum of primate lentiviruses. These investigators further showed that substitution of 375-Ser (found in most HIV-1 group M viruses) by 375-Trp (found in most SIV strains from lower primates) favored an HIV-1 Env conformation that was closer to the CD4-bound state (31–34). Based on these findings, we hypothesized that residue 375 might act as a “molecular switch” conferring enhanced Env affinity to rhesus CD4 (35) and a lower energetic barrier to conformational change following CD4 binding (31, 34, 36, 37) when the naturally-occurring Ser or Thr residues were substituted by bulky aromatic residues like Trp. In testing this hypothesis, we discovered that substitution of a single residue, 375-Ser, in primary or T/F HIV-1 Envs by Trp, Phe, Tyr, His or Met resulted in SHIVs that exhibited enhanced binding to rhesus CD4, increased infection of primary rhesus CD4^+^ T cells in culture, and consistent infection and replication by SHIVs in RMs *in vivo* (35). Importantly, these amino acid substitutions at residue 375 did not alter the tier 2 neutralization phenotype of the primary Envs nor did they appreciably alter their sensitivity to bNAbs that targeted any of the canonical bNAb recognition sites, including CD4bs, V2 apex, V3 high mannose patch or membrane proximal external region (35). Thus, it became possible, for the first time, to prospectively design SHIVs that expressed particular primary or T/F Envs, including those that elicited bNAbs in HIV-1 infected humans, and to explore parallels in the immune responses of rhesus monkeys and humans to essentially identical Env immunogens (38). This EnvΔ375 design strategy also made possible the development of SHIVs to evaluate preclinical efficacy of novel active or passive vaccination regimens against challenge by viruses bearing homologous or heterologous primary Envs (7–10). Here, we extend this work by constructing ten new SHIVs, each containing a strategically selected primary HIV-1 Env, that we then validate for retention of native antigenicity, tier 2 neutralization sensitivity and efficient replication in human and rhesus CD4^+^ T-cells *in vitro* and in RMs *in vivo*. We next describe the development and characterization of a panel of nine SHIV challenge stocks, each containing a unique tier 2 primary HIV-1 Env and grown large scale in primary rhesus CD4^+^ T-cells, for distribution as challenge strains for active or passive vaccine protection trials. We show that these SHIVs can be efficiently transmitted by different mucosal routes (rectal, vaginal, penile or oral) and that current vaccination regimens and passively administered bNAbs can prevent transmission of these viruses at neutralization titers similar to those reported in the recently concluded human Antibody-Mediated Prevention (AMP) trials (11). Finally, we describe a new second-generation design strategy that simplifies SHIV construction and eliminates extraneous *tat1* and *env* sequences, thereby making the rhesus-SHIV infection model a more readily accessible and useful research tool.

## Results

Ten primary HIV-1 Envs were chosen for SHIV constructions (**Table 1A**). These Envs were selected based on their genetic subtypes, biophysical properties, derivation from primary or T/F virus strains, and in some cases, prior development as candidate vaccine strains for human clinical trials (see **Table 1** for Env features and relevant literature citations). Env subtypes included A, B, C, AE and AG, which complement subtype A, B, C and D SHIVs that we reported previously [see (35, 38); **Table 1B**). All ten of the new SHIVs contained Envs from tier 2 viruses except for Q23.17 Env (39), which has been variably classified as tier 1b or 2 (40–42). Seven of the new SHIVs contained Envs from T/F strains of HIV-1. The 1086 Env (43) corresponds to a vaccine strain employed in the HVTN 703 efficacy vaccine trial (44–46), and the B41 Env was developed as a SOSIP trimer for potential human immunizations (47). The Ce1176 Env is from a widely used global test panel for bNAb detection (41). Env RV217.40100 is a new subtype AE T/F strain (48, 49) and Envs CH1012 and CH694 are T/F strains that elicited potent bNAbs in their respective human hosts (50, 51). Envs T250, ZM233, WITO, Q23.17 and CAP256SU were shown previously to bind unmutated common ancestors (UCAs) of human V2 apex targeted bNAbs (52–54). Thus, the Envs selected for new SHIV constructions exhibited unique pedigrees complementary to previous SHIV designs (35, 38, 55–64) that made them desirable for downstream investigations related to HIV-1 transmission, prevention, immunopathogenesis or cure.

**Table 1A.** Genetic and biological features of HIV-1 Envs used for new SHIV constructions

| HIV-1 Env | Subtype | Env GenBank Accession# | SHIV GenBank Accession# | Env properties | References |
| --- | --- | --- | --- | --- | --- |
| Q23.17 | A | AF004885 | MW410736 | Cloned from primary isolate; binds V1V2 bNAb UCAs | 39, 52-54 |
| WITO4160 | B | FJ496176 | MW410737 | T/F<sup>#</sup>; binds V1V2 bNAb UCAs | 52-54, 73 |
| B41* | B | EU576114 | MW410732 | T/F; SOSIP immunogen | 47, 73 |
| CE1176 | C | FJ444437 | MW410733 | T/F; global neutralization panel | 41, 43 |
| CH1012 | C | MG898887 | MW410734 | T/F; elicited bNAbs in human | 50 |
| ZM233 | C | DQ388517 | MW410738 | Cloned from primary isolate; binds V1V2 bNAb UCAs | 52-54, 64 |
| 1086 | C | FJ444395 | MW410739 | T/F; P5 vaccine trial | 43-46 |
| CH0694 | C | KJ700458 | MW410741 | T/F; elicited bNAbs in human | 50,51 |
| RV217.40100 | AE | MN792076 | MW410740 | T/F; Thai AE subtype | 48, 49 |
| T250-4<sup>&</sup> | AG | pending | MW410735 | Primary isolate; binds V1V2 bNAb UCAs | 20, 52-54 |

**Table 1B.**
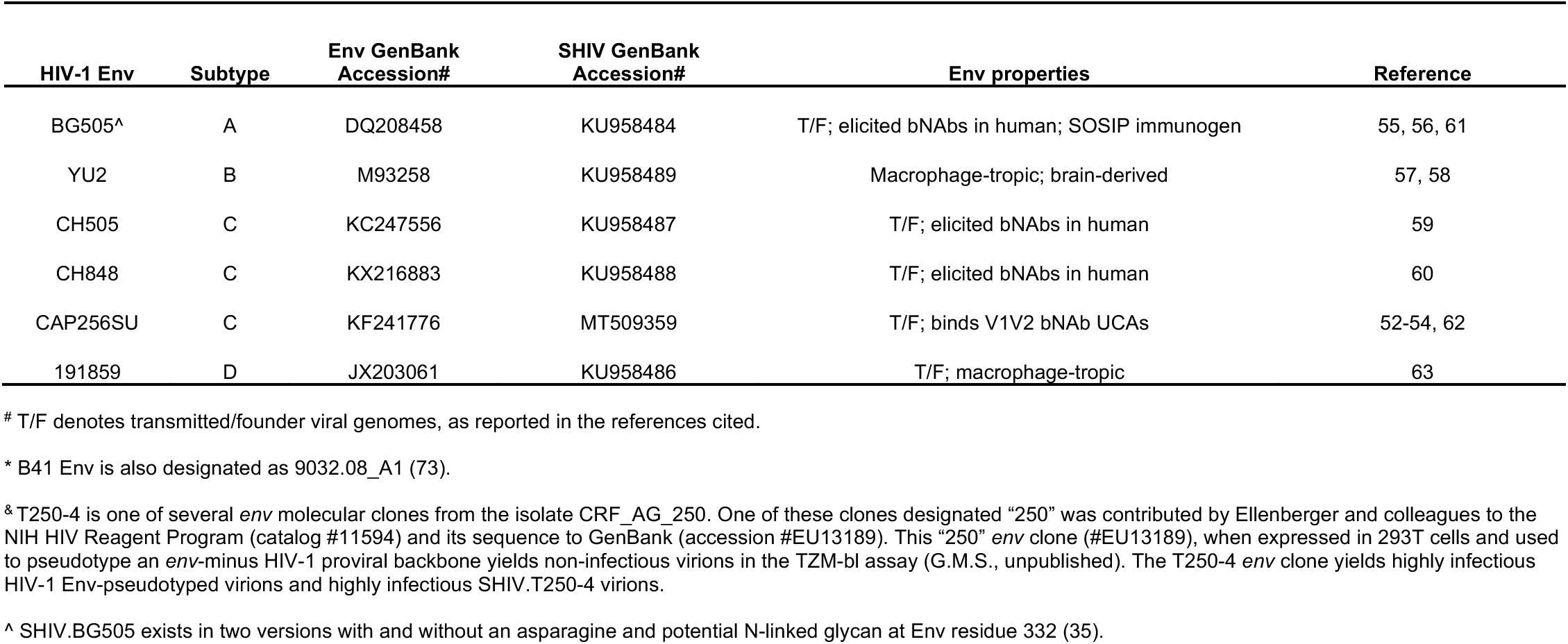
Genetic and biological features of HIV-1 Envs used in previous SHIV constructions

| HIV-1 Env | Subtype | Env GenBank Accession# | SHIV GenBank Accession# | Env properties | Reference |
| --- | --- | --- | --- | --- | --- |
| BG505<sup>^</sup> | A | DQ208458 | KU958484 | T/F; elicited bNAbs in human; SOSIP immunogen | 55, 56, 61 |
| YU2 | B | M93258 | KU958489 | Macrophage-tropic; brain-derived | 57, 58 |
| CH505 | C | KC247556 | KU958487 | T/F; elicited bNAbs in human | 59 |
| CH848 | C | KX216883 | KU958488 | T/F; elicited bNAbs in human | 60 |
| CAP256SU | C | KF241776 | MT509359 | T/F; binds V1V2 bNAb UCAs | 52-54, 62 |
| 191859 | D | JX203061 | KU958486 | T/F; macrophage-tropic | 63 |

<sup>#</sup> T/F denotes transmitted/founder viral genomes, as reported in the references cited.
\* B41 Env is also designated as 9032.08\_A1 (73).
<sup>&</sup> T250-4 is one of several *env* molecular clones from the isolate CRF\_AG\_250. One of these clones designated "250" was contributed by Ellenberger and colleagues to the NIH HIV Reagent Program (catalog #11594) and its sequence to GenBank (accession #EU13189). This "250" *env* clone (#EU13189), when expressed in 293T cells and used to pseudotype an *env*-minus HIV-1 proviral backbone yields non-infectious virions in the TZM-bl assay (G.M.S., unpublished). The T250-4 *env* clone yields highly infectious HIV-1 Env-pseudotyped virions and highly infectious SHIV.T250-4 virions.
<sup>^</sup> SHIV.BG505 exists in two versions with and without an asparagine and potential N-linked glycan at Env residue 332 (35).

The design strategy for constructing SHIVs is illustrated in **Fig. 1A**. This construction scheme allowed for the complete extracellular gp140 region of Env plus the transmembrane segment and 9 aa of the cytoplasmic tail (nucleotides 1-2153; HXB2 numbering) to be PCR-amplified *en bloc* and subcloned into a chimeric T/F SIVmac766-HIV-1 proviral backbone (35). If sequences were available for *vpu* in the source material, then the homologous *vpu-env* gp140 gene segment was amplified and subcloned into the proviral vector, since homologous *vpu-env* sequences could potentially enhance the efficiency of Env translation. Env 375 codon substitutions corresponding to Trp, Phe, Tyr, His or Met were introduced by site-directed mutagenesis into each SHIV construct, which was then prepared as a large-scale DNA stock and sequence confirmed. Genome sequences for all SHIVs were contributed to GenBank (**Table 1**). For each of the ten primary HIV-1 Envs, six variants containing the different Env375 alleles were made bringing the total number of newly constructed SHIVs to 60. In the course of SHIV constructions, we noted that certain aspects of the design scheme were inefficient, especially the requirement for multiple PCR amplifications and ligations (see Methods). We also found in SHIV infected RMs that redundant HIV-1 *tat1* and *env* gp41 sequences of 68 and 21 bp in length, respectively, that were generated as a consequence of the original cloning strategy underwent spontaneous deletion [**Figs. 1A** and S1; (35, 38)]. Thus, we modified the SIVmac766 backbone vector and amplification primers to simplify the PCR amplification step and eliminate the redundant sequences (**Fig. 1B**). We used this new design strategy to reclone SHIV.CH505, in order to perform a head-to-head comparison of viruses expressed from this new vector compared with the original SHIV design, and to clone a new SHIV containing the HIV-1 CH694 Env. Plasmid DNA for all SHIVs was transfected into 293T cells and virus-containing supernatants were characterized for p27Ag content and infectivity on TZM-bl cells. For all SHIVs, p27Ag concentrations ranged from 200-2000 ng/ml. One nanogram of p27Ag is equivalent to approximately 10^7^ virions, so SHIV titers were estimated to range from 2×10^9^ to 2×10^10^ virions per ml. We confirmed these titers by quantifying vRNA and assuming vRNA molecules per virion. Infectivity titers on TZMbl cells ranged from 2×10^5^ to 2×10^6^ per ml, corresponding to an IU to particle ratio of approximately 10^-4^. This ratio is typical for 293T-derived HIV-1 and SIV virions (35), and 100-fold lower than for virus stocks propagated in primary rhesus CD4^+^ T cells where between 1 in 100 and 1 in 50 virions are typically infectious on TZMbl cells [**Table 2** and (35)].

**Figure 1.**
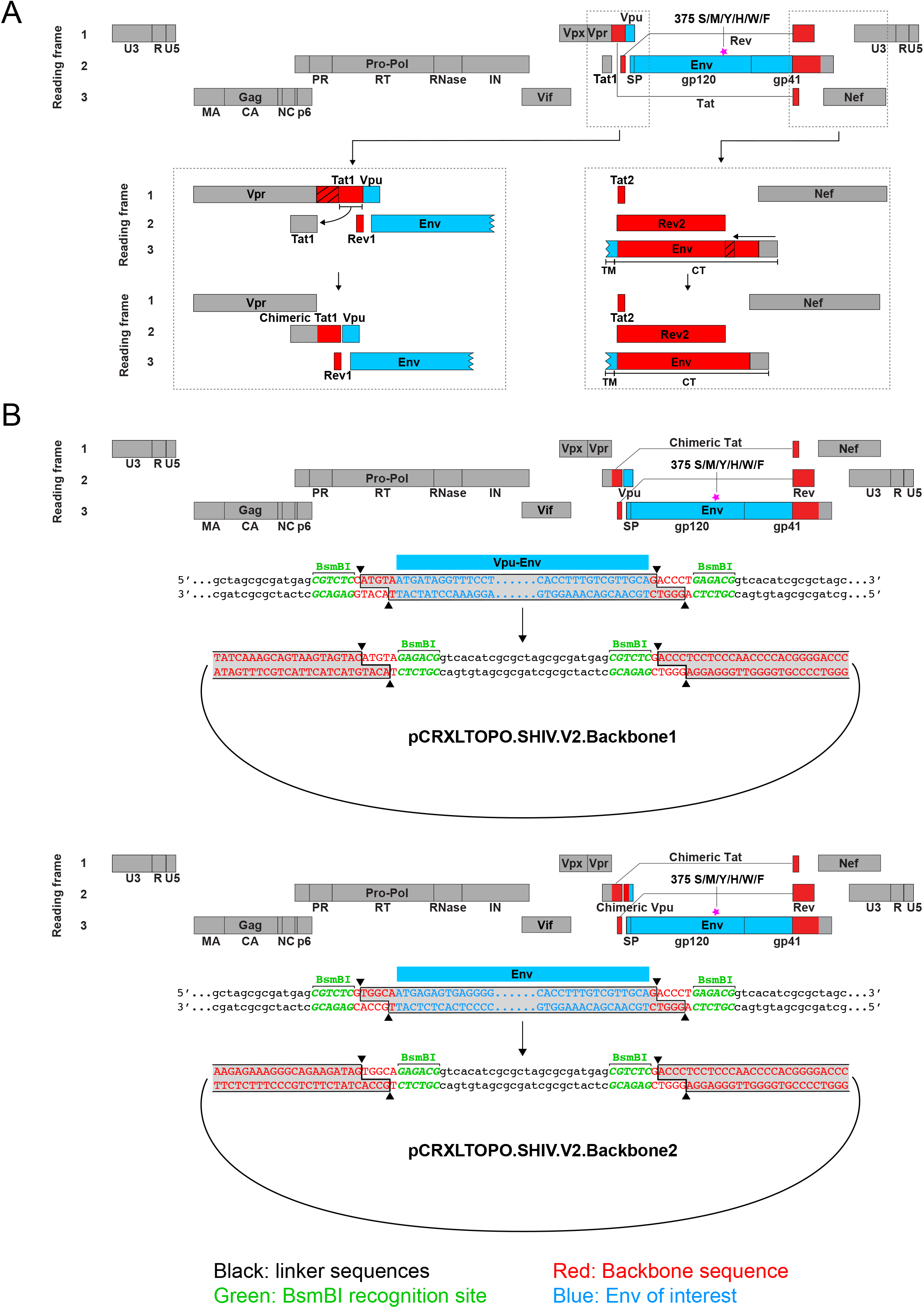
First (**A**) and second (**B**) generation design schemes for SHIV constructions. The first generation design (35) consists of a proviral backbone of SIVmac766 (a T/F clone derived from the SIVmac251 isolate) shown in grey and HIV-1.D.191859 shown in red. A *vpu-env* amplicon is subcloned into this plasmid vector as describe (35). Note in the expanded figures in panel A that the proviral backbone design (GenBank accession #KU958487) results in duplications of SIV and HIV-1 *tat1* and gp41 sequences (indicated by slash marks) that are spontaneously deleted during *in vivo* replication of the SHIVs (**Fig. S1**). The second generation design scheme (panel B) eliminates these duplications and adds a *BSMBI* cloning site that allows for simple introduction of either a *vpu-env* amplicon or *env* amplicon (GenBank accession number pending).

**Table 2.** SHIV challenge stocks expanded in primary rhesus CD4^+^ T cells

| Virus Stock | Volume/<br>vial (ml) | Date<br>generated | # vials | p27<br>ng/ml | vRNA<br>copies/ml | Infectivity Titer (IU/ml) |  | IU/Particle<br>ratio<sup>#</sup> | AID<sub>50</sub><sup>^</sup> |
| --- | --- | --- | --- | --- | --- | --- | --- | --- | --- |
|  |  |  |  |  |  | TZMbl | rhCD4<sup>+</sup> T<br>cells |  |  |
| SHIV.BG505.332N.375Y.dCT(s1<sup>▪</sup>) | 0.75 | 2/24/16 | 1,567 | 154 | 2,105,956,647 | 13,437,500 | 630,957 | 0.0128 | 1:3*, 1:120<sup>^</sup> |
|  |  |  |  | 270<sup>§</sup> | 2,263,660,000<sup>§</sup> | 6,093,000<sup>§</sup> | 165,000<sup>§</sup> | 0.0054<sup>§</sup> |  |
| SHIV.BG505.332N.375Y.dCT(s2<sup>□</sup>) | 1.0 | 1/28/19 | 2,080 | 365 | 4,089,870,000 | 31,640,000 | 4,050,000 | 0.0155 |  |
| SHIV.CH505.375H.dCT(s1) | 0.5 | 12/22/15 | 1,194 | 178 | 631,771,853 | 6,797,000 | 186,209 | 0.0215 | 1:2*, 1:80<sup>^</sup> |
| SHIV.CH505.375H.dCT(s2) | 1.0 | 5/10/17 | 1,626 | 190 | 778,255,000 | 5,234,400 | 631,000 | 0.0135 | 1:80<sup>^</sup> |
| SHIV.CH848.375H.dCT | 1.0 | 4/28/16 | 1,355 | 73 | 900,447,863 | 4,421,000 | 162,000 | 0.0098 |  |
| SHIV.191859.375M.dCT | 0.25 | 10/21/14 | 192 | 212 | 3,004,673,792 | 31,898,389 | 630,957 | 0.0212 | 1:3* |
| SHIV.B41.375H.dCT | 1.0 | 10/27/17 | 1,675 | 200 | 1,710,815,153 | 8,125,000 | 369,000 | 0.0095 |  |
| SHIV.1086.375W.dCT | 1.0 | 2/12/18 | 2,057 | 207 | 502,180,000 | 146,000 | 1,870 | 0.0006 |  |
| SHIV.CH1012.375Y.dCT | 1.0 | 2/3/20 | 2,216 | 552 | 1,576,865,000 | 4,687,000 | 125,000 | 0.0059 |  |
| SHIV.Ce1176.375HFW.dCT<sup>&</sup> | 1.0 | 2/4/20 | 2,224 | 390 | 1,619,830,000 | 6,562,000 | 125,000 | 0.0081 |  |
| SHIV.T250-4.375HWY.dCT<sup>%</sup> | 1.0 | 2/14/20 | 1,231 | 634 | 1,939,331,667 | 7,656,000 | 125,000 | 0.0079 |  |

<sup>#</sup> IU/particle ratio determined on TZMbl cells based on vRNA measurements
<sup>^</sup> AID<sub>50</sub> - Inoculum dose leading to productive clinical infection of 50% of rhesus macaques
\* Intravaginal (IVAG) inoculation route (1 ml of 1:X dilution)
<sup>^</sup> Intrarectal (IR) inoculation route (1 ml of 1:X dilution)
<sup>▪</sup> First stock expansion
<sup>□</sup> Second stock expansion
<sup>§</sup> Repeat measurements performed on samples stored for three years in vapor phase liquid nitrogen
<sup>&</sup> Stock comprised of a mixture of Env375 His, Phe and Trp variants
<sup>%</sup> Stock comprised of a mixture of Env375 His, Trp and Tyr variants

For SHIVs bearing the 10 new HIV-1 Envs, we evaluated the replication efficiency of each of them containing six different Env375 residues in primary activated human and rhesus CD4^+^ T cells *in vitro* (**Fig. 2**). With the exception of SHIV.AE.40100, which naturally contains the positively charged, aromatic residue Env375-His, none of the SHIVs containing wild-type Ser or Thr residues at position Env375 replicated appreciably in rhesus CD4^+^ T cells (**Fig. 2**). Conversely, all 10 SHIVs with wild-type Env375 residues replicated efficiently in primary activated human CD4^+^ T cells. This latter result – efficient replication of SHIVs containing wildtype 375 alleles in human CD4^+^ T cells – was an expected finding but was nonetheless critical to demonstrate, since it confirmed that the chimeric SHIVs that we made were capable of supporting replication. We next asked if substitution of the wildtype Env375 allele by one or more aliphatic or aromatic residues (Met, Trp, Phe, Tyr or His) would support SHIV replication in rhesus CD4^+^ T cells. The answer was affirmative for SHIVs expressing each of the ten HIV-1 Envs (**Fig. 2**). The differences in virus replication in rhesus CD4^+^ T cells between SHIVs expressing wild-type Env375 residues and those expressing bulky aromatic residues was generally quite large, oftentimes resulting in >100-fold differences in p27Ag concentration in culture supernatants at multiple time points throughout the infection (**Fig. 2**). Among the six different Env375 alleles that were tested, Env375-Trp most consistently supported SHIV replication in rhesus CD4^+^ T cells: it was effective in all 10 HIV-1 Env backbones. Env375-Tyr was the second most favored residue followed by Env375-His or -Phe. It is notable that Trp is also the most conserved Env375 allele across the broad evolutionary spectrum of primate lentiviruses excluding humans and great apes (30). These results thus corroborate and extend a substantial body of scientific literature indicating that SHIVs bearing primary (non-adapted) wildtype HIV-1 Envs rarely replicate efficiently in rhesus cells (1-3, 27-29, 35, 38, 65, 66) and that this restriction can be lifted by substituting a single amino acid at position Env375. In our combined studies [this manuscript plus (35, 38)], we replaced wildtype Env375 residues in 16 primary HIV-1 Envs – 15 of which could not support SHIV replication in RMs – and found in all instances that this substitution alone led to efficient SHIV replication in rhesus animals.

**Figure 2.**
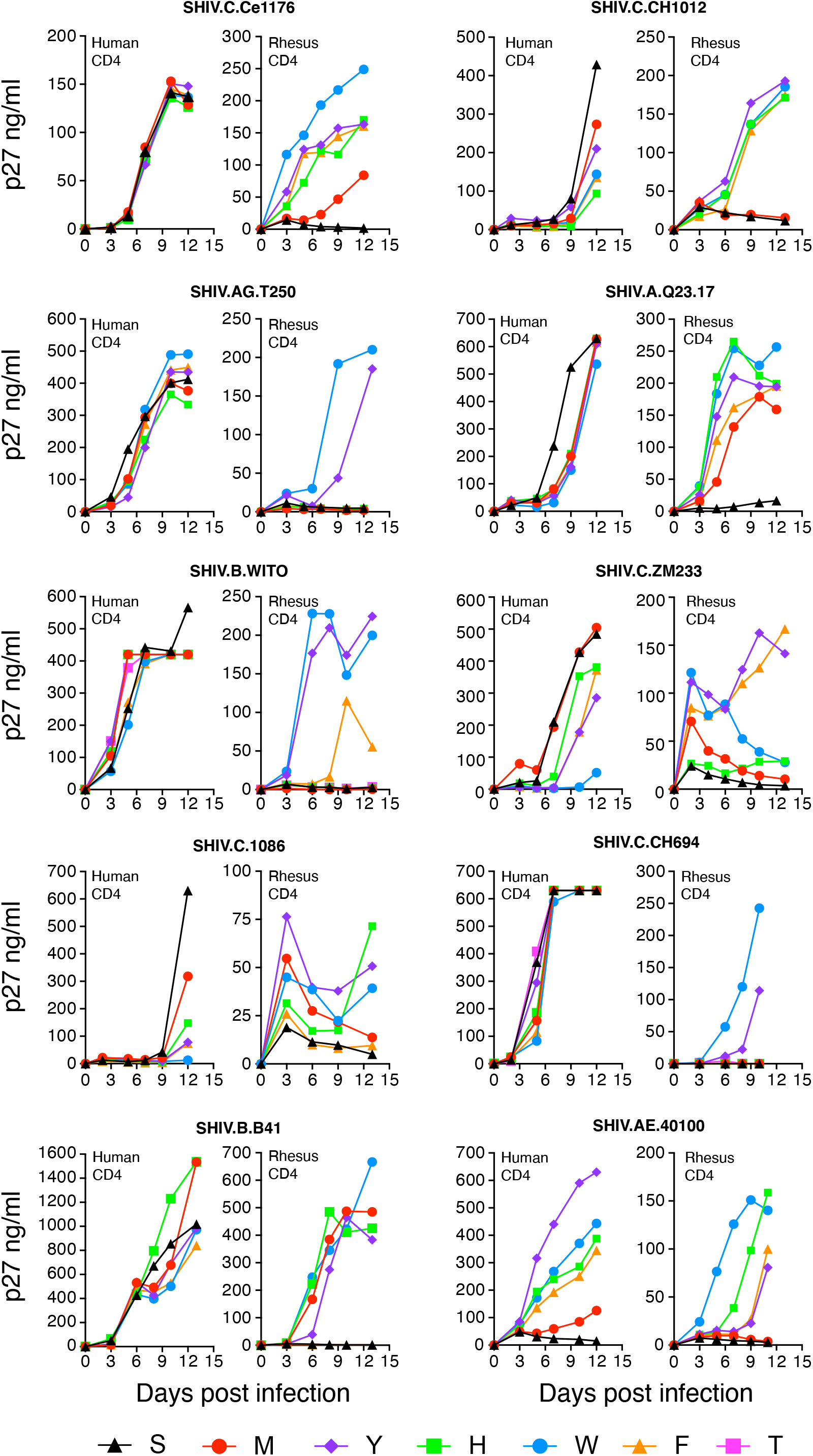
Replication kinetics of SHIVs bearing ten different HIV-1 Envs with allelic variants at residue Env375 (S-Ser, M-Met, Y-Tyr, H-His, W-Trp, F-Phe, T-Thr) in cell culture. Primary, activated human and rhesus CD4^+^ T cells were inoculated with 293T-derived SHIVs and culture supernatants were sampled on the days indicated. Panels display results of representative experiments, which were reproduced in large scale expansions of SHIVs in rhesus CD4^+^ T cells *in vitro* (**Table 2**) and in RMs *in vivo* (Fig. 3).

To extend these findings to *in vivo* analyses, we inoculated 41 RMs intravenously in groups of 3 to 6 animals each, with SHIVs containing one of the ten selected HIV-1 Envs and an equal mixture of the six Env375 alleles (**Table S1** and **Fig. 3**). We used this experimental design for two reasons: First, because target cell availability is not limited in the initial two weeks of infection during which time virus titers increase exponentially (67–70), we could use deep sequencing of plasma vRNA/cDNA to directly compare the relative replication rates of the six Env375 allelic variants in an *in vivo* competitive setting. Second, it would be impractical and prohibitively expensive to test 60 SHIVs individually in 60 different monkeys, and even if this could be done, the results would be confounded by monkey-specific variables such as MHC class I and II recognition. Each of the 41 RMs that we inoculated with a SHIV Env375 mixture became productively infected after a single challenge (**Fig. 3**). In most animals, peak viremia occurred at day 14 post-SHIV inoculation and plasma virus load setpoints were reached 16-24 weeks later. Animals treated with anti-CD8 mAb at the time of SHIV inoculation developed significantly higher peak and setpoint viremia titers compared with untreated animals (p<0.01 for both). A subset of animals was treated with anti-CD8 mAb at setpoint, 20-50 weeks after infection; most of these animals exhibited increases in virus titers. We performed next generation sequencing (NGS) on plasma samples taken 2 and 4 weeks post-infection to determine the relative replication rates of the different Env375 allelic variants (**Fig. 3**). We expected that differences in infectivity of the Env375 variants would be reflected in the plasma virus quasispecies by two weeks post-inoculation since the combined half-lives of circulating virus and the cells producing it is <1 day (71), resulting in multiple rounds of *de novo* virus infection and replication during this early interval. This was indeed the case. Overall, there was a good correlation between Env375 residues that supported SHIV replication *in vitro* and *in vivo*. For example, in all ten different Env backgrounds, Env375-Ser failed to support SHIV replication in primary rhesus CD4^+^ T cells *in vitro* (**Fig. 2**) and the same was true in RMs *in vivo* (**Fig. 3**). Conversely, Env375-Trp supported SHIV replication in all ten Env backgrounds *in vitro* and was a predominant allele supporting efficient SHIV replication in 7 of 10 Env backgrounds *in vivo*. There were some differences in Env 375 residues that best supported SHIV replication *in vitro* versus *in vivo*. For example, for SHIVs bearing ZM233 and CH0694 Envs, 375-Trp supported efficient virus replication *in vitro* but not *in vivo*, where 375-Tyr was dominant. And the Env375-His allele, which is naturally present in most subtype AE viruses including the AE.40100 strain, supported efficient SHIV.AE.40100 replication in rhesus CD4^+^ T cells *in vitro* but not *in* vivo. Taken together, the findings indicate that substitution of wildtype Env375 alleles in primary HIV-1 Envs with Trp, Tyr or His results in SHIV chimeras that replicate efficiently in RMs. However, since it is impossible to predict with certainty which Env375 allele will best support *in vivo* replication of a SHIV bearing any particular HIV-1 Env, an *in vivo* competition experiment similar to that illustrated in **Fig. 3** must be conducted.

**Figure 3.**
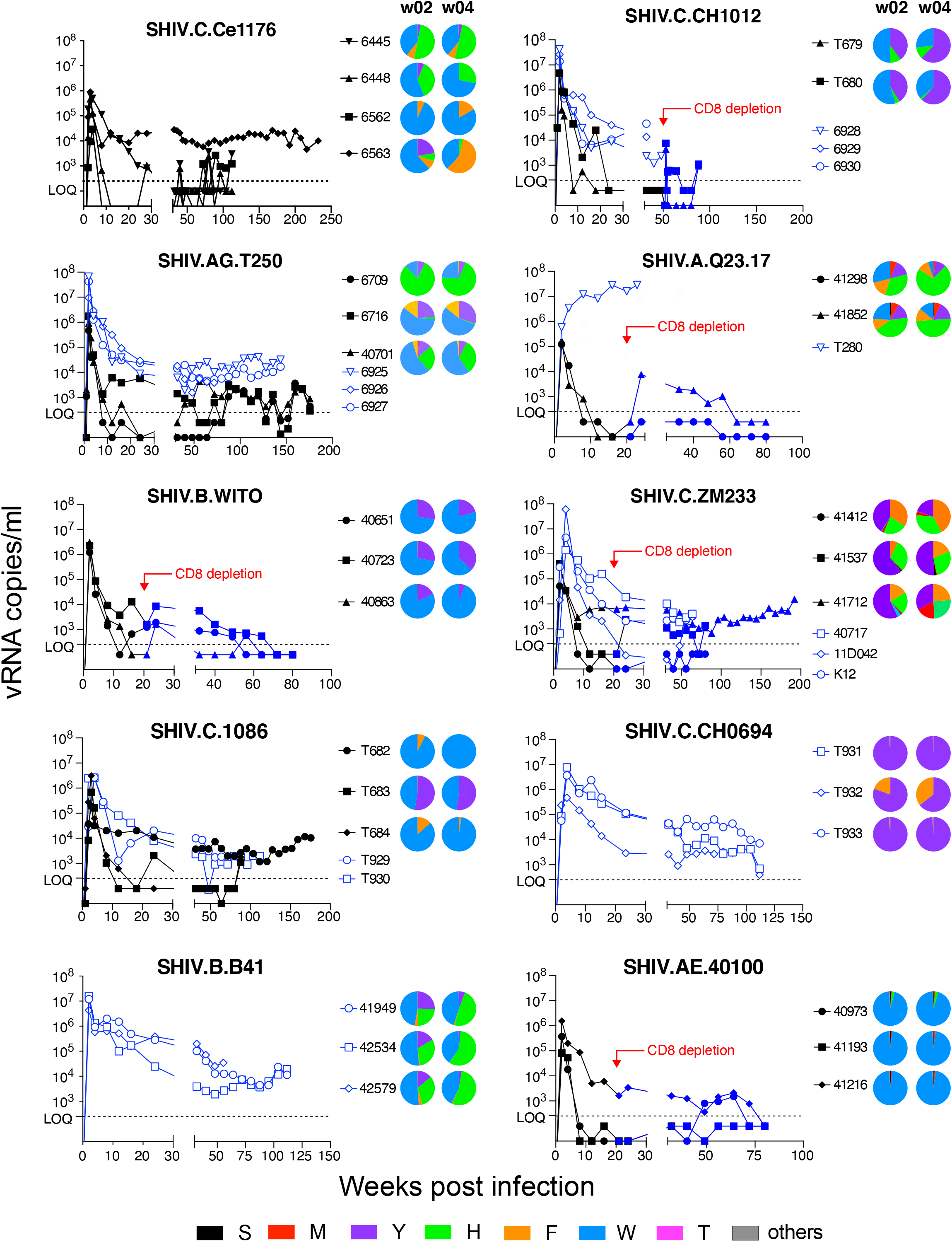
Plasma vRNA kinetics following intravenous inoculation of RMs with SHIVs bearing six Env 375 allelic variants. Open symbols denote animals that were treated with anti-CD8 mAb at week 0. Solid black symbols denote animals that were not treated with anti-CD8 mAb at week 0. Some animals were treated with anti-CD8 mAb at time points indicated and these animals are indicated by a shift in solid symbols from black to blue. Pie diagrams represent >5,000 vRNA/cDNA sequences and indicate the proportions of different Env375 alleles in plasma vRNA 2 and 4 weeks after SHIV inoculations. Where pie diagrams are not shown, animals were inoculated with SHIVs containing a single Env375 allele (**Table S1**).

We also compared the relative replication efficiency of SHIV.CH505.375H generated by the first and second generation construction strategies (**Fig. S2**). We showed previously that in animals infected by viruses produced from the first generation design, that redundant HIV-1 *tat1* and *env* gp41 sequences (68 and 21 bp, respectively) were spontaneously deleted following prolonged *in vivo* replication [**Fig. S1**; (35, 38)]. This suggested a fitness disadvantage for viruses containing the redundant sequences, leading us to hypothesize that animals infected by an equal mixture of the viruses derived from the two designs would show preferential replication by viruses lacking the redundant sequences. This was indeed the case (**Figs. S1A** and **S1B**). At three weeks post-infection, viruses lacking the redundant sequences comprised >95% of the plasma virus quasispecies, and by week 8, they comprised >99% of plasma virus.

To be a relevant model for HIV-1 vaccine studies, SHIV Envs should exhibit clinically relevant antigenic profiles, neutralization sensitivity phenotypes, and coreceptor usage indistinguishable from the primary HIV-1 Envs from which they were derived. We evaluated the neutralization sensitivity patterns of Envs expressing the wild-type Env375 allele compared with Envs expressing one or more of the alternative Env375 alleles that were found to support replication in rhesus CD4^+^ T cells *in vitro* (**Figs. 2**) and in RMs *in vivo* (**Figs. 3**). SHIVs were analyzed using polyclonal anti-HIV-1 sera and a battery of monoclonal antibodies (mAbs) that bind canonical bNAb epitopes, linear V3 epitopes or CD4-induced (CD4i) epitopes (**Fig. 4**). Linear V3 and CD4i epitopes are generally concealed on native Env trimers from primary viruses (40, 72–74), and thus mAbs targeting these epitopes typically fail to neutralize primary virus strains. Conversely, neutralization by linear V3 or CD4i mAbs is generally an indication of a non-native “open” trimer structure typical of laboratory-adapted viruses. In none of the ten primary Env backbones that we tested did Env375 substitutions result in neutralization by linear V3 or CD4i mAbs (**Fig. 4**). Nor did Env375 mutations alter the neutralization sensitivity of these Envs to HIVIG B, HIVIG C or a high titer, broadly neutralizing HIV-1 infected patient plasma specimen CH1754. These results suggest that the Envs bearing residue 375 substitutions retained their native or near-native conformation. These Envs also retained their antigenicity with respect to bNAb epitope presentation since mAbs targeting CD4bs, V2 apex, V3 high mannose patch, and MPER sites exhibited similar neutralization patterns against wild-type and Env375 substituted variants. It is notable that the contours of the neutralization curves, the IC_50_, IC_80_ and IC_90_ values, and the steep sigmoidal inflections were generally indistinguishable between wildtype Envs and Envs bearing residue 375 substitutions. SHIV.Q23.17 demonstrated neutralization sensitivity patterns to the bNAb mAbs, the three polyclonal anti-HIV IgG and plasma reagents and the mAbs targeting linear V3 or CD4i epitopes that were similar to the other nine SHIVs, thus supporting a tier 2 status for this virus. We also tested SHIV.CH505.375H derived by first and second generation design schemes for sensitivity to HIV-1 bNAbs, linear V3 targeted mAbs, HIVIG-C and the anti-HIV-1 broadly neutralizing polyclonal plasma CH1754: the two virus preparations showed indistinguishable neutralization sensitivity patterns (**Fig. S2**). Finally, the SHIVs containing the ten new HIV-1 Envs were tested for coreceptor usage by analyzing their sensitivity to AMD-3100 (a CXCR4 inhibitor) and Maraviroc (a CCR5 inhibitor). Maraviroc, but not AMD-3100, inhibited the entry of all 10 SHIVs in the TZM-bl entry assay (**Fig. S3**), thus demonstrating CCR5-dependent entry. Altogether, the results indicate that Env375 substitutions did not appreciably alter the antigenicity, tier 2 neutralization sensitivity or CCR5 tropism of any of the ten SHIVs.

**Figure 4.**
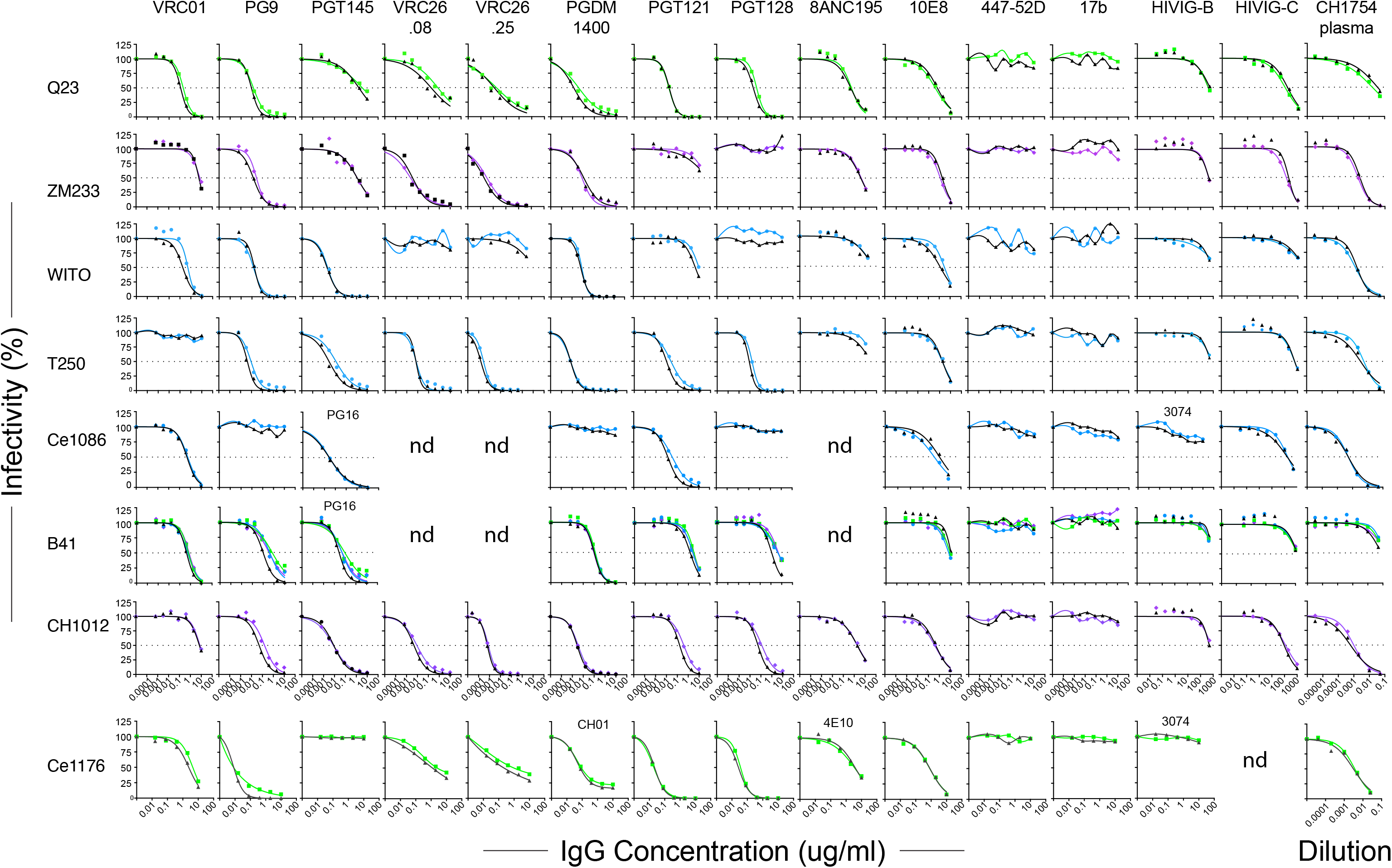
Neutralization sensitivity of SHIVs bearing wildtype Env375 residues (black symbols) or alternate residues preferred for replication in RMs (Trp – blue, His – green, Tyr – purple). 293T cell derived virus was assayed for entry into TZMbl cells in the presence or absence of anti-HIV-1 mAbs, IgG preparations (HIVIG-B, -C), or human plasma CH1754 (103).

SHIVs intended for use as challenge strains in preclinical vaccine trials can be generated from 293T cells by transfection of proviral DNA or by virus passage and expansion in primary human or rhesus CD4^+^ T cells. Each approach has potential advantages and disadvantages (3, 75). We chose to prepare challenge stocks by infecting primary, activated rhesus CD4^+^ T cells with molecularly cloned virus derived from 293T cell transfections and then expanding the virus as rapidly as possible so as to minimize chances for culture adaptation. By this means, we could ensure that the viral envelopes of challenge stocks contained exclusively rhesus (not human) membrane-associated proteins and that glycosylation patterns would be of rhesus (not human) origin. We selected nine SHIV strains for large scale expansion in rhesus cells and these are listed in **Table 2**. These SHIVs were chosen to be representative of global HIV-1 diversity, including subtypes A, B, C, D, and AG, and to include SHIVs bearing BG505.N332, CH505 and 1086 Envs, which correspond to vaccine candidates in current or recent human clinical trials. Our aim was to generate large numbers of identical replicates of each SHIV stock (>1,000 vials per SHIV), which could then be characterized biophysically for genetic composition, particle content, infectivity, antigenicity and neutralization sensitivity and cryopreserved in vapor phase liquid nitrogen (<160°C) for subsequent distribution as validated, standardized SHIV challenge stocks. Thus, we inoculated cultures of 100-200 million primary, activated, rhesus CD4^+^ cells pooled from three naïve Indian RMs at a multiplicity of infection (MOI) of approximately 0.01 with genetically homogeneous, sequence-confirmed, 293T transfection-derived virus stocks. For SHIV.Ce1176, we infected primary rhesus cells with an equal mixture of Env375-His, Phe and Trp alleles, and for SHIV.T250 we infected cells with an equal mixture of Env375-His, Tyr and Trp alleles, because each of these alleles in these two Env backgrounds had shown preferential replication in different animals (**Fig. 3**). The other SHIV challenge stocks were generated with single Env375 alleles (**Table 2**). On days 7 and 14 post-SHIV inoculation, we added new media and approximately 100-200 million fresh, uninfected rhesus CD4^+^ T cells from three different naïve RMs so as to expand cell numbers and culture volumes while maintaining cell concentrations between 1-2 million per milliliter. Beginning on day ∼10 post-SHIV inoculation, we collected the total volume of culture supernatant and replaced it with a greater volume of fresh medium. This complete media collection and replacement was then repeated every 4 days through day 21. By this means, we could collect as much as 2.5 liters of culture medium containing each SHIV over a period of approximately 21 days. Each supernatant collection was centrifuged twice at 2500 rpm (1000g) for 15 minutes to remove any residual cells or cell debris and then immediately frozen in bulk at −80°C. Supernatants were not filtered so as to retain the highest possible infectivity titers. Thus, most of the virus that was collected and frozen during the 18-21 day culture period was <4 days old and underwent only one freeze-thaw cycle prior to final vialing. After all supernatant collections had been made, they were thawed at room temperature, combined in a sterile 3 liter flask to ensure complete mixing, and then aliquoted into as many as 2,500 cryovials, generally at 1 ml per vial. The vials were then transferred to vapor phase liquid nitrogen for long-term storage. By this means, we could ensure that all vials were virtually identical in their contents. Between 192 and 2,224 vials per SHIV, each containing between 0.25 and 1.0 ml of challenge stock, were cryopreserved (**Table 2**). Validation analyses were done on thawed cryovial samples to ensure results would be representative of all cryopreserved samples. Challenge stocks were free of bacterial or fungal contamination based on culture on thioglycolate broth. p27Ag concentrations ranged from 73 to 634 ng/ml and vRNA concentrations ranged from 5.0 x 10^8^ to 4.1 x 10^9^ vRNA/ml. Infectivity was tested on TZM-bl cells where it ranged from 1.5 x 10^5^ to 3.2 x 10^7^ IU/ml, and on primary rhesus CD4^+^ T cells where it ranged between 1.9 x 10^3^ to 4.1 x 10^6^ IU/ml. The genetic composition of the SHIV challenge stocks was analyzed by single genome sequencing of 3’ half-genomes to validate the authenticity of each stock and to determine if there was evidence of selection *in vitro* (**Fig. 5A**). Stocks of SHIV.Ce1176 and SHIV.T250 were sequenced by Illumina deep sequencing to determine the relative proportion of the different Env375 alleles in the final challenge stocks (**Fig. 5B**). Envelope sequence mean and maximum diversity averaged 0.05% (range 0.03-0.13%) and 0.30% (range 0.15-0.42%), respectively in the nine challenge stocks. Mutations across the complete gp160 were essentially random in all challenge stocks except in a secondary expansion of SHIV.CH505. This challenge stock was prepared by infecting naïve rhesus CD4^+^ T cells with virus from the first expansion of SHIV.CH505 in an attempt to expand sequence diversity and increase infectivity titers. Maximum sequence diversity and maximum sequence divergence from the T/F sequence were 0.35% and 0.29% for stock #2 compared with 0.15% and 0.08%, respectively, for stock #1. p27Ag and vRNA concentrations and infectivity titers on TZM cells were similar for stocks #1 and #2 and infectivity titers on primary rhesus CD4 T cells were about 3-fold higher for stock #2 compared with stock #1.

**Figure 5.**
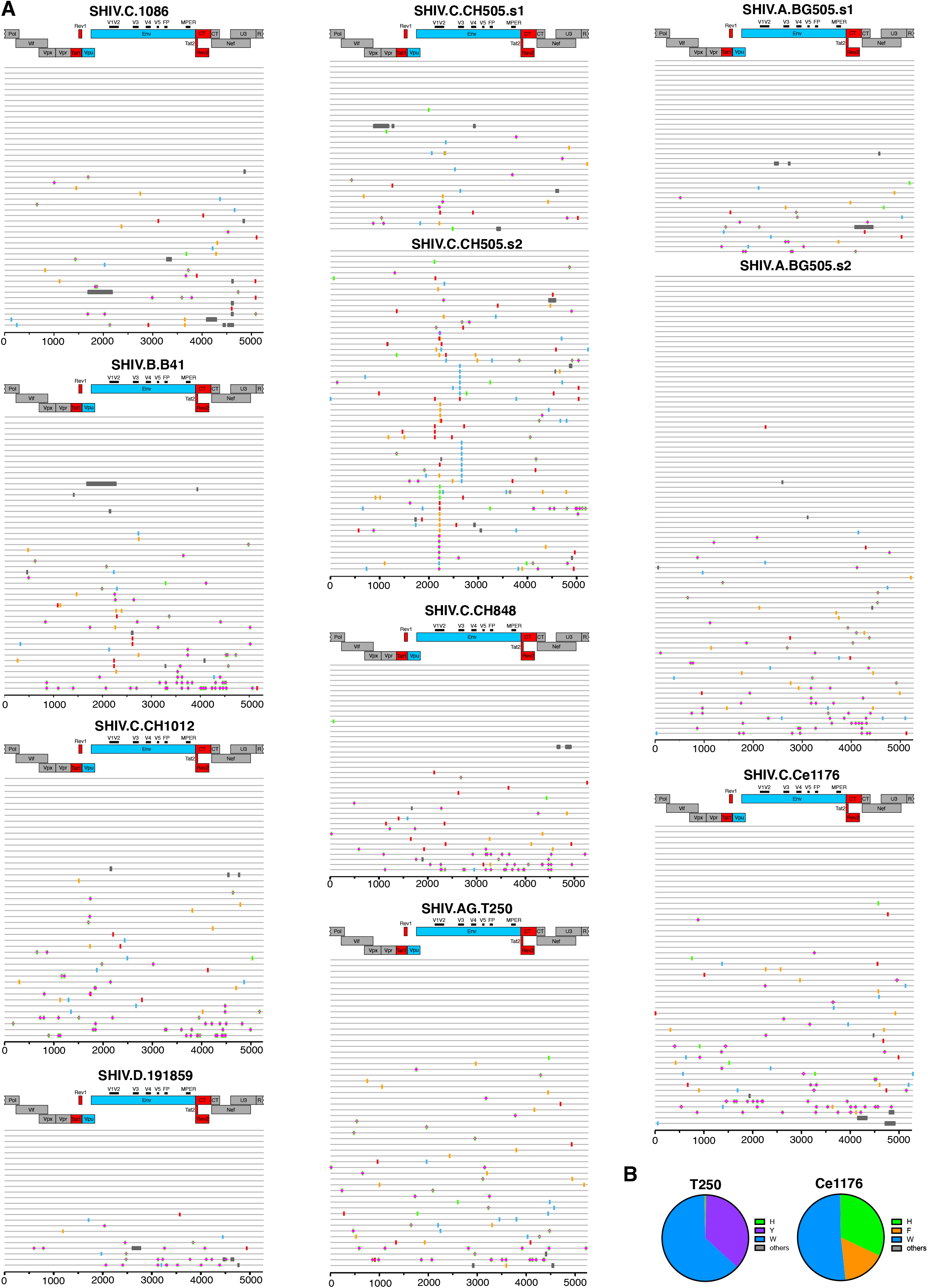
(**A**) Pixel plots (https://www.hiv.lanl.gov/content/sequence/pixel/pixel.html) of single genome sequences of 3’ half genomes of rhesus CD4+ T cell grown SHIV challenge stocks. Tic marks indicate nucleotide differences from the SHIV molecular clones (A – red, C – yellow, T – blue, G – green, APOBEC site – green+pink, INDELs – grey). (**B**) Pie diagrams showing the relative proportions of different Env375 alleles in SHIV.C.1176 and SHIV.AG.T250 challenge stocks (see text for explanation).

HIV-1 strains produced in primary human CD4^+^ T cells, compared with the same viruses produced in 293T cells, have been reported to exhibit variably greater resistance to neutralizing antibodies (76, 77). These differences have been attributed to differences in Env content, cell adhesion molecules, surface glycan composition or other factors (75). We tested five SHIVs – BG505, CH505, CH848, B41, D.191859 – produced in primary rhesus CD4^+^ T cells and in 293T cells for sensitivity to 19 neutralizing mAbs that targeted CD4bs, V3 glycan, V2 apex, MPER, surface glycan, CD4i or linear V3 epitopes (**Fig. 6**). None of the viruses, regardless of cell derivation, were sensitive to the four mAbs that targeted CD4i or linear V3 epitopes, indicating that they retained a native-like closed Env trimer regardless of the cell of origin. SHIVs produced in 293T cells and primary rhesus cells also exhibited similar overall patterns of sensitivity to the other 15 mAbs in that if a SHIV was sensitive (or resistant) to neutralization by a particular mAb, then this was true regardless of its cell of origin. However, as reported for HIV-1 strains, we observed enhanced resistance to some mAbs by some SHIVs grown in primary rhesus cells compared with 293T cells. This difference was two-to five-fold for all five SHIV strains when exposed to VRC01 and 3BNC117 and as much as 25-fold for certain other virus-antibody combinations such as BG505-CH01, CH505-CH01, CH505-PGT145, CH505-VRC26.25, B41-CH01 and B41-VRC26.25 (**Fig. 6**).

**Figure 6.**
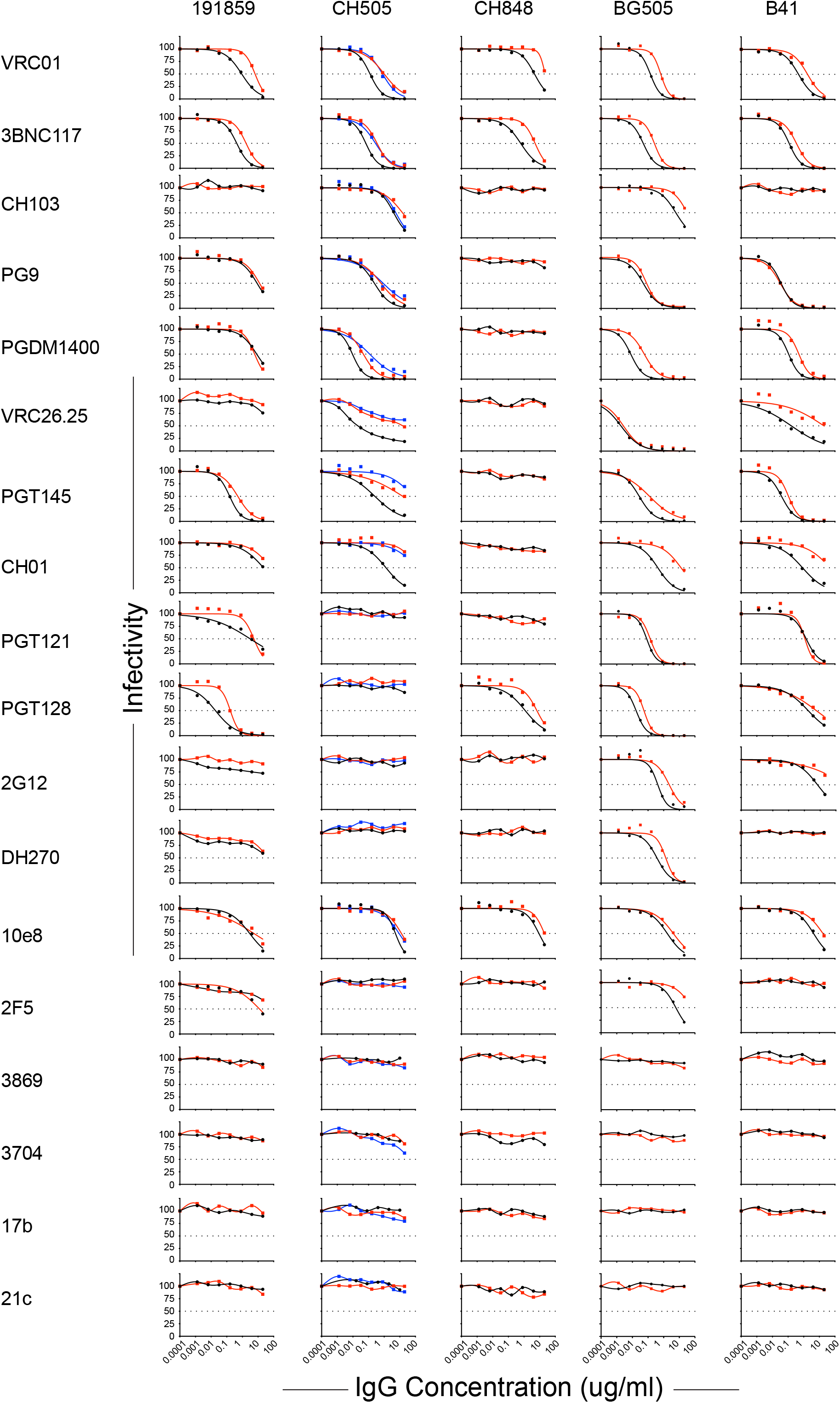
Neutralization sensitivity of SHIVs bearing rhesus-preferred Env375 residues and generated in either 293T cells (black symbols) or primary rhesus CD4^+^ T cells (red symbols). For SHIV.C.CH505, the rhesus CD4^+^ T cell derived stock 1 is depicted by red symbols and stock 2 by blue symbols.

The properties of rhesus CD4^+^ T cell grown SHIV challenge stocks summarized in **Table 2**, especially their consistently high virus titers and infectivity measurements, suggested that these virus strains might be suitable for mucosal transmission studies and to assess the preclinical efficacy of actively-induced or passively-administered bNAbs. Nearly all natural routes of HIV-1 acquisition result from transmission across mucosal surfaces, the exceptions being intrauterine and intravenous infections. Previously, we showed that SHIVs BG505, CH505 and D.191859 can be transmitted efficiently across rectal, vaginal and oral mucosae (35), resulting in productive clinical infection with virus replication kinetics and plasma virus titers indistinguishable from human infections by HIV-1 (69, 70). Penile acquisition is another important route of HIV-1 transmission in humans (78), and **Fig. 7A** shows that SHIV.D.191859 can be transmitted by atraumatic inoculation of foreskin and glans. Peak viremia occurred at approximately 2 weeks post-challenge and plasma virus load setpoint was reached by 6 weeks. Setpoint viremia persisted at 50,000 to 200,000 vRNA molecules per milliliter through 16 weeks of follow-up when the experiment was terminated per protocol. These kinetics of SHIV.D.191859 replication post-penile transmission were similar to plasma virus load kinetics of the same SHIV strain transmitted by intrarectal, intravaginal and intravenous routes (**Fig. 7A**).

**Figure 7.**
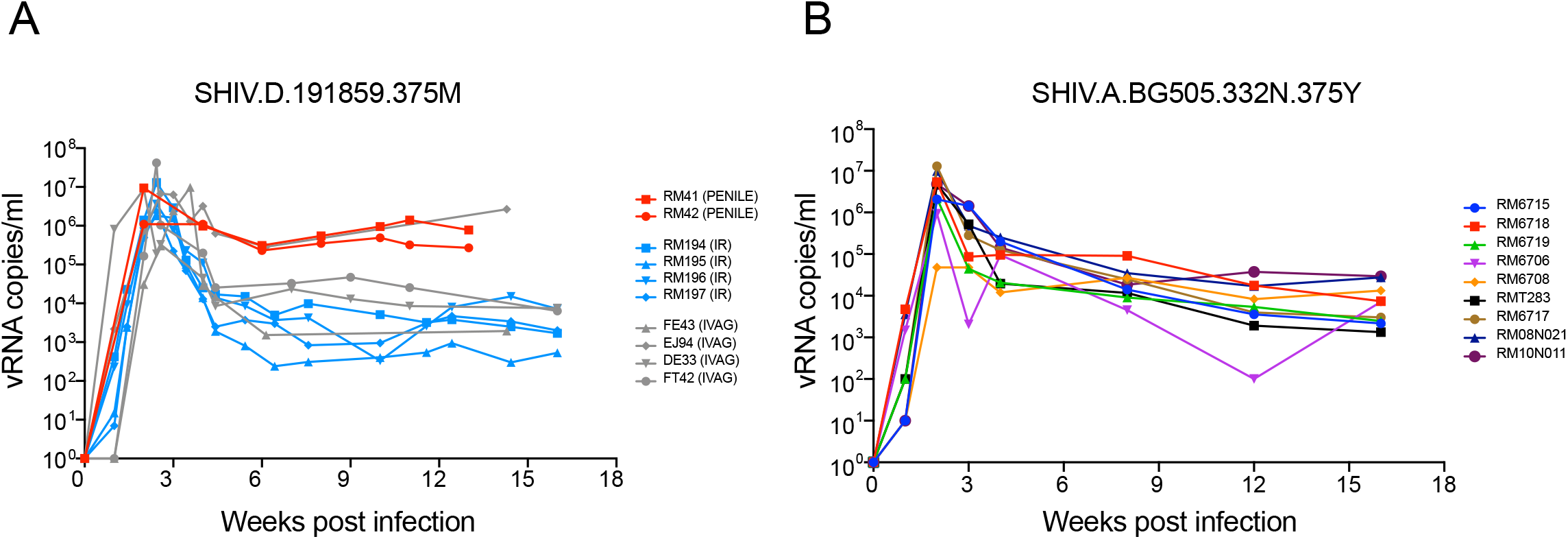
Plasma viral load kinetics following atraumatic penile inoculation of SHIV.D.191859 (**A**) and atraumatic intrarectal inoculation of SHIV.A.BG505.N332.375Y (**B**). Two RMs (RM41 and RM42) were inoculated atraumatically a single time with 0.25 ml of undiluted SHIV.D.191859 stock into the sulcus between penile glans and foreskin. This resulted in productive clinical infection in both animals, as indicated by plasma viremia one week later. Plasma viral load kinetics through 13 weeks of followup were comparable to SHIV.D.191859 plasma virus loads resulting from other routes of transmission reported earlier (35, 101) and similar to plasma viremia of SHIV.A.BG505.N332.375Y following low-dose intrarectal challenge (**B**).

Lastly, we performed a low-dose, repetitive challenge rectal titration of SHIV.BG505.N332 in 12 naïve RMs to estimate the AID_50_ of the challenge stock and to assess plasma viral load kinetics following IR infection. Three of 4 inoculations at a dose of 1:20 (1 ml), 3 of 4 inoculations at a dose of 1:100 (1 ml), and 3 of 9 inoculations at a dose of 1:160 (1 ml) resulted in productive clinical infection. Acute and early SHIV.BG505.N332 replication kinetics (**Fig. 7B**) were similar to mucosal infection by SHIV.D.191859 (**Fig. 7A**) and also similar to the ten SHIVs illustrated in **Fig. 3** that were infected intravenously. Although our intrarectal AID_50_ titration experiment for SHIV.BG505.N332 involved a small number of animals (n=12) and was subject to stochastic effects related to intrarectal virus inoculation, we could nonetheless estimate the AID_50_ of this stock to be approximately 1:120 (1 ml) for atraumatic IR challenge. This result was corroborated in the control (sham-treated) arm of a preclinical trial assessing the protective efficacy of BG505 SOSIP vaccine elicited neutralizing antibodies against a homologous SHIV.BG505.N332 challenge (7).

## Discussion

In recent years, there have been notable advances in HIV prevention and cure research (79–83) yet the goals of effective vaccination and cure – even a “functional” cure – seem far in the distance. Increasingly, experimental medicine trials in humans have been pursued as a strategy to accelerate translational research (82), but at the same time, there remain untapped opportunities and needs for animal models to complement and synergize with human studies to hasten progress. Different scientific questions demand different model systems, ranging from transgenic or humanized mice to outbred small and large animals. Aside from the great apes, which are endangered and thus precluded from laboratory investigation, the rhesus macaque monkey (*Macaca mulatta*) is most similar to humans in its immune repertoire (84, 85). For HIV-related investigations in primates, two classes of viruses are broadly used: simian immunodeficiency viruses (SIV) and chimeric SHIVs that express HIV-1 Envs within an SIV background (3). The present study adds 10 new SHIVs to the research portfolio of HIV investigators; characterizes key biological properties of these SHIVs that are relevant to virus transmission, prevention, immunopathogenesis and cure research; and describes a new SHIV design strategy and cloning vector that can facilitate future SHIV constructions.

The HIV-1 Env glycoprotein is critical to virus transmission, persistence and pathogenesis since it conveys the essential functions of receptor and coreceptor binding and membrane fusion. At the same time, Env is the target of an array of neutralizing antibodies and cytotoxic T-cells that cause it to evolve continuously in order to escape recognition that would otherwise lead to virus elimination (38, 86, 87). Env accomplishes the latter by means of highly evolved properties, including occlusion of trimer-interface epitopes (88), epitope variation (89), conformational masking (90) and glycan shielding (91). Although HIV-1 Env is notorious for its variability and global diversity (www.hiv.lanl.gov), it is nonetheless constrained in its potential for immediate or near-term evolution due to the myriad of essential biological functions encoded in its sequence (38, 92–94). These constraints can be lifted, however, by *in vitro* cultivation (66, 95) or extensive passage in unnatural animal hosts (1–3, 22). The implication of these observations is that the most relevant HIV-1 Envs for studies of vaccine-elicited protection, passively acquired antibody protection, or curative intervention are primary or T/F Envs from viruses that are responsible for clinical transmission and the establishment of persistent infection in humans (7–10, 96). T/F Envs express the precise primary, secondary, tertiary and quaternary protein structures that are essential for transmission and T/F Envs are the ones that a vaccine-elicited bNAb response must recognize if it is to be protective (38, 73, 82). Envs derived from short-term virus cultures in human lymphocytes or Env sequences derived from plasma vRNA/cDNA are a first approximation to T/F Envs but they may differ in important but unrecognized features. Envs derived from extensively passaged virus cultures are less likely to reflect the biologic and antigenic properties of T/F viruses. In this context, 7 of the 10 new SHIVs described in the current study, and 12 of 16 SHIVs that we have reported overall (**Table 1**), were constructed using T/F Envs. The remainder was constructed using primary Envs.

A recent study by Keele and colleagues (29) aimed to create new subtype C T/F SHIVs using 20 South African subtype C T/F Envs and either of two strategies to enhance replication in primary RM CD4^+^ T cells. One of these strategies was the same EnvΔ375 design employed here and the other was an EnvΔ281 approach reported elsewhere (28). Because the O’Brien study pooled SHIVs for competitive replication analyses in RMs, a precise determination of the proportion of wild-type HIV-1 Envs that could support SHIV replication in monkeys could not be made. However, in the instances where EnvΔ375 substitutions were made and the resulting SHIVs were tested individually, EnvΔ375 substitutions were successful in conferring replication competence to SHIVs in rhesus cells. The addition of EnvΔ281 was neither additive nor synergistic. In our studies described here (**Table 1**) and elsewhere (35, 38), we created a total of 16 EnvΔ375 SHIVs, and each one replicated efficiently in RM CD4+ T cells *in vitro* and in RMs *in vivo*. Altogether, the results suggest that EnvΔ375 substitution is an effective means for creating SHIVs that have a high likelihood of replicating efficiently in RMs. The second-generation design strategy illustrated in **Figs. 1B** and **S1** can facilitate this process by substantially reducing the time and effort required to construct new SHIVs and by improving their replication fitness.

A useful outbred primate model for HIV-1 infection of humans should be rational in design, amenable to iterative changes in the challenge viruses, and consistent in reproducing relevant features of disease. Previously, SHIV infections of RMs did not always meet these requirements since SHIVs replicated variably in RMs and often required *in vitro* or *in vivo* adaptations to achieve consistent infection or replication. Oftentimes, these changes were not fully understood mechanistically nor were their immunobiological effects fully appreciated. Moreover, *in vitro* measures of virus content, infectivity and replication in cell culture did not always predict *in vivo* outcomes, lending a measure of uncertainty to SHIV design and analysis. The EnvΔ375 strategy alleviates much of this uncertainty and unpredictability as demonstrated by the following results: i) Env375 substitutions alone were sufficient to enhance Env affinity to rhesus CD4, reduce the energetic threshold for downstream Env transitions following CD4 binding, and convey efficient infectivity to the virus in primary rhesus CD4^+^ T cells *in vitro* and *in vivo*; ii) the Env375 substitution strategy worked consistently; every attempt that we (**Table 1**), Keele (29) and Barouch (97) have made to engineer a T/F or primary HIV-1 Env SHIV by residue 375 substitution has succeeded in producing a chimeric virus that replicates efficiently in RMs; iii) the ability of such EnvΔ375 SHIVs to replicate *in vivo* was, in each case, predicted by efficient replication in primary, activated rhesus CD4^+^ T cells *in vitro*; this is a different result from what has been reported for other SHIVs (1–3, 65) and we suspect that our simple EnvΔ375 design scheme, our protocol for rhCD4+ T cell activation, and our method for infecting these cells in tissue culture are responsible for the differences; iv) the antigenicity and tier 2 neutralization sensitivity of wildtype HIV-1 Envs was closely mirrored by EnvΔ375 mutants expressed from 293T cells or as infectious SHIVs from primary rhesus CD4 T cells; v) the genetic diversity of each SHIV infection stock was very low when virus was expressed either from 293T cells or from primary rhesus CD4^+^ T-cells; vi) transmission efficiency of SHIVs across rhesus rectal, vaginal, penile and oral mucosa, and intravenously, mirrored the transmission efficiency of HIV-1 in humans; vii) acute and early SHIV replication dynamics in RMs measured by plasma vRNA replicated what has been seen in humans, including a 7-14 day eclipse period before vRNA is detectable in plasma, an exponential increase in plasma virus load to a peak approximately 14-28 days post-infection, establishment of setpoint viremia two or more months later, and immunopathogenesis leading to clinically-defined AIDS in a subset of animals (69, 70, 73); viii) SHIV infected RMs consistently elicited autologous, strain-specific NAbs, and in some cases bNAbs, with kinetics similar to HIV-1 infected humans (35, 38); ix) molecular pathways of SHIV Env evolution in RMs closely mirrored evolution of homologous HIV-1 Envs in humans, including precise molecular patterns of Env-Ab coevolution leading to Nab escape, and in some animals, the development of bNAbs (38). The latter results speak to the native-like structure of SHIV Envs and to homologies and orthologies in human and rhesus immunoglobulin gene repertoires (38, 85). Altogether, the findings highlight the reproducibility and relevance of the SHIV EnvΔ375 infected RM as a model system for HIV-1 infection in humans.

Efficient mucosal transmission leading to productive clinical infection with consistent patterns of plasma viremia are critical features of SHIVs intended for use as challenge strains to test for vaccine efficacy and for mechanistic studies of virus transmission. We tested SHIVs BG505.N332, CH505, D.191859 and T250-4 for mucosal transmission and titered each challenge stock for 50% animal infectious doses (AID_50_). For these studies, virus challenge stocks were grown in primary rhesus CD4^+^ T cells. Challenge stocks were first subjected to thorough analytical measurements of virus concentration, infectivity, genotypic complexity and phenotype with respect to coreceptor usage and antigenicity (**Table 2** and **Figs. 5, 6, S3**). Because of the wide scientific interest of BG505.N332 SOSIP as a vaccine candidate, we conducted a low-dose atraumatic intrarectal titration study of SHIV.BG505.N332 (**Fig. 7B**) where we estimated the IR AID_50_ of this stock to be approximately 1:120 (1 ml). Burton and colleagues (7) corroborated this estimate by showing that 6 of 6 naïve RMs inoculated intrarectally with a 1:20 (1 ml) dose of this same challenge stock, and 9 of 12 naïve RMs inoculated intrarectally with a 1:75 (1 ml) dose, became productively infected after a single challenge (7). Importantly, these results demonstrated reproducibility in clinical infectivity titers of the identical challenge stock used at different primate centers and in animals obtained from different breeding colonies. Replication dynamics of SHIV.BG505.N332 following the low dose intrarectal inoculations were quite similar in the two studies [**Fib. 7B** and (7)]: a meta-analysis of the results revealed peak viremia geometric mean titers of 2.7 x 10^6^ vRNA/ml at day 14 post-challenge and plasma viral load setpoint geometric mean titers of 9.2 x 10^3^ vRNA/ml by week 12, with 23 of 24 animals remaining viremic. Pulendran and colleagues used this same SHIV.BG505.N332 challenge stock for low dose intravaginal challenges in a preclinical protection study in RMs (9). In a control arm of 15 sham vaccinated RMs, they found the AID_50_ to be approximately 1:3 (1 ml). Peak plasma viremia (GMT = 1.7 x 10^6^ vRNA/ml) was again at 14 days post-infection and plasma viral load setpoint was reached by week 10, with 14 of 15 animals remaining viremic (GMT = 1.7 x 10^3^ vRNA/ml). The 40-fold difference in AID_50_ between IR and IVAG challenge routes is consistent with previous findings with SHIV and SIVs (3, 98, 99) and is similar to estimates of relative infectivity in humans exposed to receptive anal intercourse versus receptive vaginal intercourse (78). We also titrated SHIV.CH505 challenge stocks for AID_50_ in RMs following intrarectal or intravaginal inoculation. In independent studies with a total of 21 RMs, Klatt [(100) and unpublished data] and Haynes estimated the AID_50_ following IR challenge of naïve RMs to be approximately 1:80 (1 ml), while Felber and colleagues (8) found the AID_50_ of this stock following IVAG challenge to be approximately 1:2 (1 ml) (**Table 2**). These findings again demonstrate reproducibility in AID_50_ titers in different primate centers and in monkeys from different breeding colonies as well as a 30-40 fold difference in infectivity between IR versus IVAG challenge routes. Previously, we estimated the AID_50_ for SHIV.D.191879 for IVAG inoculation to be approximately 1:3 (1 ml) (101). Here, we could not estimate an AID_50_ for penile transmission by the SHIV.D.191879 challenge stock since 2 of 2 animals became infected after a single inoculation (**Fig. 7A**), but the findings suggest that the AID_50_ titers of this stock for penile transmission are likely to be sufficient for it to be used as a challenge stock in preclinical prevention trials once formal titering is completed. Finally, in an ongoing study, Sok, Rakasz and colleagues have estimated the AID_50_ of SHIV.T250 to be approximately 1:160 (1 ml inoculum) following atraumatic rectal inoculation (unpublished). Thus, in multiple studies of mucosal infection by BG505.N332, CH505, D.191859 and T250, AID_50_ titers and post-infection plasma viral load kinetics were consistent between SHIVs and between studies conducted at different primate facilities and mirrored analytical assessments of the different challenge stocks *in vitro* (**Table 2**). These findings suggest that precise measurements of virion content and infectivity of different challenge stocks correlate well with AID_50_ titers following mucosal challenge, which is important because it can facilitate AID_50_ titrations of new challenge stocks going forward.

Altogether, the findings of this study suggest that the SHIVs listed in **Table 1** can be broadly useful as challenge stocks for preclinical studies of vaccine-elicited or passively-acquired antibody protection; for assessing novel cure interventions; and for mechanistic studies of virus transmission and pathogenesis. We have contributed the rhCD4 T cell grown SHIV challenge stocks and the 16 SHIV plasmid DNA stocks to the NIH NIAID HIV Reagent Repository and to the Penn Center for AIDS Research Virology Core Laboratory, which provides investigators with derivative reagents (e.g., barcoded SHIVs for lineage-tracing, sequence-verified viral DNA maxipreps, minimally-adapted T/F SHIV variants with enhanced *in vivo* replication dynamics, titered 293T derived infectious virus stocks) to meet research needs. One important research application that we anticipate in the future is in comparative efficacy testing of different vaccines against the same heterologous tier 2 primary virus challenge stock, and the same vaccine against a single heterologous virus administered by different mucosal inoculation routes. Such studies promise to inform HIV-1 immunogen design and testing as new vaccine candidates are developed.

## Acknowledgments

We thank the staff at Bioqual, Inc., for exceptional care and assistance with experiments involving nonhuman primates, and T. Demarco, N. DeNaeyer and members of the Nonhuman Primate Virology Core Laboratory at the Duke Human Vaccine Institute for SHIV plasma vRNA measurements. We thank N. Miller, J. Warren, Q. Dang and S. Staprans for helpful discussions regarding the development of SHIVs as a research community resource. This work was funded by the Bill & Melinda Gates Foundation (OPP1145046, OPP1206647/INV-007939) and the Division of AIDS (NIAID/NIH) through support of the Duke Center for HIV/AIDS Vaccine Immunology-Immunogen Discovery (UM1 AI100645) and Consortium for HIV/AIDS Vaccine Development (UM1 AI144371), the Penn Center for AIDS Research (P30 AI045008), and grants AI088564, AI094604, AI131251, AI131331, AI050529, and AI150590. R. Roark was supported by an NIH training grant in HIV Pathogenesis (T32-AI007632). This project has been funded in part with Federal funds from the National Cancer Institute, National Institutes of Health, under Contract No. HHSN261200800001E and 75N91019D00024. The content of this publication does not necessarily reflect the views or policies of the Department of Health and Human Services, nor does mention of trade names, commercial products, or organizations imply endorsement by the U.S. Government.

## Materials and Methods

### Nonhuman primate care and procedures

Indian rhesus macaques were housed and studied at Bioqual, Inc., Rockville, MD, or at the Plum Borough animal facility at the University of Pittsburgh, Pittsburgh, PA, according to guidelines and standards of the Association for Assessment and Accreditation of Laboratory Animal Care. Experiments were approved by the Bioqual, University of Pittsburgh, Duke University and University of Pennsylvania Institutional Animal Care and Use Committees. All animals were negative for Mamu-A*01, B*08, and B*17 and screened as negative for retroviruses, measles, ebola and *T. cruzi*. Animals were sedated for blood draws, anti-CD8 mAb infusions, and SHIV inoculations. For estimations of AID_50_ of SHIV challenge stocks, animals were inoculated atraumatically by penile, rectal or vaginal routes. Penile inoculations were performed in anesthetized animals in a recumbent supine position. The foreskin was retracted vertically and laterally and separated from the glans exposing the preputial mucosa and coronal sulcus. 0.25 ml of undiluted SHIV challenge stock was slowly and carefully inoculated into this potential space using a 1 ml syringe and the vertical-lateral foreskin retraction maintained for 20 minutes. Intrarectal or intravaginal inoculations were performed by inserting a flexible lubricated pediatric feeding tube atraumatically 3-5 cm into the rectum or vagina of animals lying in a Trendelenburg position followed by the administration of diluted virus stock in a total volume of 1 ml by slow push. Animals were maintained in Trendelenburg position for 30 minutes before being repositioned and returned to their cages to recover from anesthesia. Intravenous SHIV inoculations were performed by placing an intravenous catheter into the femoral vein of anesthetized animals in supine position, administering small volumes of sterile normal saline and confirming venous access by retrograde blood return. SHIVs bearing any of 10 different HIV-1 Envs, each with as many as six Env375 allelic variants in a total volume of 1 ml DMEM medium, were administered by slow push. Following virus inoculation, the IV line was flushed with normal saline. A subset of animals then received an intravenous infusion of 25 mg/kg of anti-CD8alpha mAb [MT807R1; NIH Nonhuman Primate Reagent Resource, NHPRR (https://www.nhpreagents.org/)] or anti-CD8beta mAb (CD8beta255R1; NHPRR) mAb over a period of 3-5 minutes. Another subset of animals received anti-CD8 mAb at 18-48 weeks post infection. RMs were phlebotomized weekly, then monthly, and then every other month to collect and cryopreserve blood samples.

### Processing and storage of rhesus and human blood specimens

Blood samples from rhesus macaques were collected in sterile 10 ml vacutainers containing ACD-A anticoagulant. Up to 40 ml of ACD-A anticoagulated blood from each RM was combined in a sterile 50 mL polypropylene conical tube, centrifuged at 2100 rpm (1000xg) for 10 min at 20°C, and the plasma collected in a fresh 50 mL conical tube without disturbing the buffy coat WBC layer and large red cell pellet. The plasma was centrifuged again at 3,000 rpm (∼2000xg) for 15 minutes at 20°C in order to remove all platelets and cells. Plasma was collected and aliquoted into cryovials and stored at −80°C. The RBC/WBC pellet was resuspended in an equal volume of Hanks balanced salt solution (HBSS) without Ca^++^ or Mg^++^ and containing 2mM EDTA and then divided into four 50 ml conical tubes. Additional HBSS-EDTA (2mM) buffer was added to bring the volume of the RBC/WBC mixture to 30 ml in each tube. The cell suspension was then carefully underlayered with 14 ml 96% Ficoll-Paque and centrifuged at 1800 rpm (725xg) for 20 min at 20°C in a swinging bucket tabletop centrifuge with slow acceleration and braking so as not to disrupt the ficoll-cell interface. Mononuclear cells at the ficoll interface were collected and transferred to a new 50ml centrifuge tube containing HBSS-EDTA (w/o Ca^++^ or Mg^++^) and centrifuged at 1000 rpm (∼200 g) for 15 min at 20°C. This pellets PBMCs and leaves most of the platelets in the supernatant. The supernatant was removed and the cell pellet was resuspended in 40 ml HBSS (with Mg^++^/Ca^++^ and without EDTA) + 1% FBS. To remove additional contaminating platelets, the cell suspension was centrifuged again at 1000 rpm (∼200xg) for 15 minutes at 20°C and the supernatant discarded. The cell pellet was tap-resuspended in the residual 0.1-0.3 ml of media and then brought to a volume of 10 ml HBSS (with Mg^++^/Ca^++^) plus 1% FBS. Cells were counted and viability assessed by trypan blue exclusion. Cells were centrifuged again at 1200rpm (300xg) for 10 min at 20°C, the supernatant discarded, and the cells resuspended at a concentration of 5−10×10^6^ cells/ml in CryoStor cell cryopreservation media (Sigma Cat. C2999), aliquoted into 1ml cryovials (CryoClear cryovials; Globe Scientific Inc., Cat. 3010), placed in a Corning CoolCell LX cell freezing container, stored overnite at −80°C, and then transferred to vapor phase liquid N_2_ for long-term storage. Alternatively, freshly isolated rhesus PBMCs were processed immediately for CD4^+^ T cell purification and activation. Human PBMCs from de-identified normal blood samples were isolated by similar procedures from leukopaks obtained from the University of Pennsylvania Comprehensive Cancer Center Human Immunology Core Laboratory and either cryopreserved or used immediately for CD4^+^ T cell purification and activation.

### SHIV constructions

SHIVs were constructed in one of two chimeric SIV/HIV proviral backbone plasmids. The original backbone (**Fig. 1A**) was first described by Li et al. (35) and was used in that study to generate SHIV.A.BG505, SHIV.B.YU2, SHIV.C.CH505, SHIV.C.CH848 and SHIV.D.191859. This backbone was subsequently employed by Roark to generate SHIV.C.CAP256SU (38) and by other investigators to generate still other SHIVs, all based on this EnvΔ375 design strategy (29, 97). We designated the first generation plasmid as pCRXTOPO.SHIV.v1.backbone1 and pCRXTOPO.SHIV.v1.backbone2. Version 1 backbones 1 and 2 allow for the cloning of *vpu-env* (*env* nucleotides 1-2153, HXB2 numbering) or *env*-only (*env* nucleotides 1-2153, HXB2 numbering) amplicons, respectively. This first generation plasmid required cumbersome sequential PCR amplifications and ligations in order to generate a complete replication competent chimeric SHIV provirus. In addition, the first generation design scheme generated an SIV – HIV-1 *tat1* redundancy and an SIV – HIV-1 *env* gp41 redundancy, both of which were spontaneously deleted when SHIVs replicated persistently *in vivo* [e.g., see **Fig. S1A** and (35, 38)]. We thus engineered second generation SHIV cloning vectors designated pCRXTOPO.SHIV.v2.backbone1 and pCRXTOPO.SHIV.v2.backbone2, which allow for cloning of the identical *vpu-env* and *env*-only amplicons, respectively. In the first generation SHIV backbone, unique restriction enzyme recognition sites for *BstBI* and *XhoI* are present in the middle of SIV vpx and after the 3’ LTR in the vector sequence, respectively. We synthesized two fragments that contain these two enzyme sites and the genes in between. We eliminated the redundant *tat1* and *env* gp41 sequences and replaced the *vpu-env* and *env* genes with a linker fragment that carries two *BsmBI* restriction enzyme sites (**Fig. 1**). The *BsmBI* site appended at the N-terminus of the linker recognizes the reverse complementary DNA strand and creates a 3’ overhang; the one added at the C-terminus recognizes the positive strand DNA and creates a 5’ overhang. This design results in two different sticky ends, which allows unidirectional cloning of the insert into the backbone. *BsmBI* is a Type IIS restriction enzyme that cleaves outside of its recognition site and thus the enzyme recognition sequence does not remain after ligating the insert into the backbone (**Fig. 1**). The two synthesized fragments were then cloned into original SHIV backbone separately using the BstBI and XhoI sites. The resulting two version 2 SHIV backbones were then used for cloning *vpu-env* (1-2153, HXB2) and *env* (1-2153, HXB2) gene, respectively. The *vpu*-*env* gp140 segments of HIV-1 Ce1176, CH1012, T250-4, Q23.17, WITO, ZM233, 1086, B41 and 40100 were cloned into the first generation SHIV backbone using methods described previously (Li, et al., 2016). The *vpu*-*env* gp140 segments of HIV-1 CH694 and CH505 were cloned into the second generation SHIV backbone by appending the *BsmBI* recognition sequences to 5’ and 3’ ends of the amplicon and performing a standard ligation (35). We then used the QuikChange II XL Site-Directed Mutagenesis kit (Agilent Technologies) to create allelic variants (M, Y, F, W, or H) of the wild type Env375-Ser or -Thr codons. Wild type and mutant plasmids were transformed into MAX Efficiency Stbl2 Competent Cells (Invitrogen) for maxi-DNA preparations. Each 10-kb viral genome was sequenced in its entirety to authenticate its identity and genome integrity. Infectious SHIV stocks were generated in 293T cells as previously described.

### SHIV Infection of primary rhesus and human CD4^+^ T cells

Purified rhesus and human CD4^+^ T cells were isolated from PBMCs using magnetic MACS CD4 MicroBeads (Miltenyi Biotec), as previously described (35). They were activated by incubation with anti-biotin MACSiBead particles (Miltenyi Biotec) loaded with biotinylated anti-CD2, -CD28 and -CD3 mAbs, as previously described (35). The replication kinetics of each of the SHIVs and Env375 variants in primary, activated human and rhesus CD4^+^ T cells were determined again as previously described (35). Briefly, 293T supernatants containing 300 ng p27Ag of each variant, were added to 2 x 10^6^ purified human or rhesus CD4 T cells in complete RPMI growth medium (RPMI1640 with 15% heat-inactivated fetal bovine serum (FBS, Hyclone), 100 U/mL penicillin–streptomycin (Gibco), 30 U/mL IL-2 (aldesleukin, Prometheus Laboratories) and 30 µg/ml DEAE-Dextran. 300 ng p27Ag is equal to ∼3 x 10^9^ virions, ∼3 x 10^5^ IU on TZMbl cells, or ∼3 x 10^4^ IU on primary CD4^+^ T-cells, so the estimated MOI of this titration was estimated to be between 0.01 and 0.05. The cell and virus mixtures were incubated for 2 hours under constant rotation at 37°C to facilitate infection, washed three times with RPMI1640, and resuspended in complete RPMI1640 medium lacking DEAE-Dextran. Cells were plated into 24-well plates at 2 million cells in 1 ml and cultured for 13 days, with sampling of 0.2ml supernatant and media replacement every 2-3 days. Supernatants were assayed for p27Ag concentration by ELISA according to manufacturer’s instructions (Zeptometrix).

### SHIV challenge stock generation in primary rhesus CD4^+^ T cells

100-200 million primary, activated, rhesus CD4^+^ cells pooled from three naïve RMs at a concentration of 10^7^ cells/ml in complete RPMI1640 medium with 10% FCS and DEAE-Dextran (30 ug/ml) were inoculated with 293T cell-derived SHIVs at a MOI of 0.1-0.5 in TZM-bl cells and an estimated MOI of 0.01-0.05 in primary rhesus CD4^+^ T cells. For SHIV.Ce1176, we infected primary rhesus cells with an equal mixture of Env375-His, Phe and Trp alleles, and for SHIV.T250 we infected cells with an equal mixture of Env375-His, Tyr and Trp alleles, because these alleles in these two Env backgrounds had shown differential replication in different animals (**Fig. 3**). The other SHIV challenge stocks were generated with viruses containing single rhesus-preferred Env375 alleles (**Table 2**). Total volume of the SHIV-cell mixture was typically 10-30 ml, depending of the infectivity titers of the 293T virus stock. The SHIV-cell mixture was transferred to a T75 flask, which was fixed to a rotating wheel or rocker so that leakage or spillage was not possible. This apparatus was then placed in a 37°C 5% CO_2_ incubator for 2 hours of continuous mixing. The contents of the T75 flask were then transferred to a sterile 50 ml polypropylene tube and centrifuged at room temp at 1200 rpm (∼300xg) for 10 min. Supernatant was decanted, the cells gently tap resuspended in the residual medium (<0.5 ml) and then resuspended in 50 ml complete RPMI medium with 10% FCS and the wash step repeated twice. The washing steps are important to remove DEAE-dextran, which can be toxic to cells in culture, and to remove unbound virus. Cells were then resuspended at a concentration of 1−2×10^6^ cells/ml in complete RPMI1640 medium with 10% FCS, Il-2 and antibiotics in T100 flasks and incubated at 37°C in a 5% CO_2_ incubator. On days ∼7 and ∼14 post-SHIV inoculation, additional fresh media and approximately 100-200 million fresh, uninfected, activated rhesus CD4^+^ T cells from three different naïve RMs were added to the cultures, which were transferred into T250 flasks to accommodate larger volumes. This expansion of the cultures markedly increased cell numbers and supernatant volumes while maintaining cell concentrations between 2-4 million per milliliter. The culture supernatant was sampled on approximately days 1, 4, 7, 10, 14, 17 and 20 for p27Ag concentration with assays performed weekly. Typically, p27Ag concentrations were <50 ng/ml on day 7 but rose rapidly to >200 ng/ml by day 10. On day ∼10 post-SHIV inoculation, the total volume of culture supernatant was collected, centrifuged twice at 2500 rpm (1000xg) for 15 minutes to remove any residual cells or cell debris, and then frozen in bulk at −80°C. The supernatant was replaced with a greater volume of fresh medium as additional uninfected activated rhesus CD4^+^ T cells were added and as cells divided, again keeping cell concentrations a 2-4 million per milliliter. Between days 10 and 21, p27Ag production was maximal and concentrations in the supernatant rose rapidly to >200 ng/ml every 3-4 days after each complete collection of culture supernatant. By this means, we could collect as much as 2.5 liters of culture medium containing each SHIV over a three week culture period. Importantly, because complete supernatant collections and fresh media replacements were performed every 3-4 days beginning on day ∼10 post-SHIV inoculation, most of the virus that was collected and frozen was <4 days old and underwent only one freeze-thaw cycle prior to final vialing. Once all supernatant collections had been made over the 18-21 day culture period, they were thawed at room temperature at the same time, combined in a sterile 3 liter flask to ensure complete mixing, and aliquoted into as many as 2,500 cryovials, generally at 1 ml per vial. The vials were then transferred to a −80°C freezer overnight and then to vapor phase liquid nitrogen for long-term storage. By this means, we could ensure that all vials were essentially identical in their contents.

### Virus entry and neutralizing antibody assays

Assays for virus entry and neutralizing antibodies were performed using TZM-bl indicator cells, as previously described (35, 91). The NAb assay is essentially identical to that employed by Montefiori, Seaman and colleagues (102) (https://www.hiv.lanl.gov/content/nab-reference-strains/html/home.htm), the only difference being that we plate virus and test plasma or mAbs or purified polyclonal IgG onto adherent TZM-bl cells and hold the concentration of human and rhesus plasma/serum constant across all wells at 10%. In addition to this 10% final concentration of plasma/serum, the culture medium consists of Dulbecco’s Modified Eagle’s Medium (DMEM) with 40 *μ*gm/ml of DEAE-dextran and pen/strep antibiotics. Infections were performed in duplicate. Uninfected cells were used to correct for background luciferase activity. The infectivity of each virus without antibodies was set at 100%. The 50% inhibitory concentration (IC_50_) is the antibody concentration that reduces by 50% the RLU compared with the no Ab control wells after correction for background. Nonlinear regression curves were determined and IC_50_ values calculated by using variable slope (four parameters) function in Prism software (v8.0). In the virus entry assay used to determine infectivity titers of 293T cell-derived viruses (**Table 2**), the culture medium consists of DMEM with 10% FBS, 40 *μ*gm/ml DEAE dextran and pen/strep antibiotics and cell entry is quantified by beta-galactosidase expression after 48 hours, as described (35).

### Coreceptor use analysis

TZM-bl cells were seeded in 96-well plate at density of 15,000 cells per well and cultured overnight at 37°C with humidified air and 5% CO_2_. Cells were incubated with selective entry inhibitors for 1 hour, followed by inoculation of 2,000 TZMbl IU of virus per well. Coreceptor inhibitors included 10 μM Maraviroc (CCR5), 1.2 μM AMD3100 (CXCR4), a mixture of inhibitors or media control. Viral Envs YU2 (CCR5-tropic) and SG3 (CXCR4-tropic) were included as controls. The infectivity of the media control wells was set at 100%. The infectivity of the experimental wells was quantified by percent of infection compared with the media control wells after correction for background.

### Plasma viral RNA quantification

Plasma viral load measurements were performed by the AIDS and Cancer Virus Program, Leidos Biomedical Research Inc., Frederick National Laboratory and by the NIH/NIAID-sponsored Nonhuman Primate Virology Core Laboratory at the Duke Human Vaccine Institute, as previously described (35, 38). Over the course of this study, the sensitivity limits for accurate vRNA quantification using 0.5 ml of NHP plasma improved from 250 RNA cp/ml to <100 RNA cp/mL. We chose a conservative threshold of 100 RNA cp/mL for a limit of detection and 250 RNA cp/mL for the limit of quantification.

### Viral RNA Sequencing

Single genome sequencing of SHIV 3’ half genomes was performed as previously described (35, 73). Geneious software was used for alignments and sequence analysis. The sequences were visualized using the LANL Highlighter plot tools (https://www.hiv.lanl.gov/content/sequence/HIGHLIGHT/highlighter_top.html). To analyze the prevalence of 375 variants, next-generation sequencing was performed using Illumina MiSeq system as described (35, 38). For each animal, 20,000-200,000 vRNA copies were used for reverse transcription and bulk RT-PCR. Raw reads from each bulk PCR were analyzed and the frequency of S, T, M, Y, H, W, and F codons at position 375 was determined by using Geneious software.

### Statistical analyses

Statistical tests were calculated by using GraphPad Prism 8 software. The Mann-Whitney test was used to determine whether the peak and setpoint viral loads of anti-CD8 treated animals were significantly different from untreated animals. We chose a nonparametric rank-based test because both peak and setpoint viral loads of the untreated group failed the D’Agostino & Pearson normality test (P-values < 0.05). The geometric means were calculated using the Column statistics function of GraphPad Prism 8. The mean and maximum diversities were calculated using Poisson-Fitter v2 program (https://www.hiv.lanl.gov).

---

**Figure S1.**
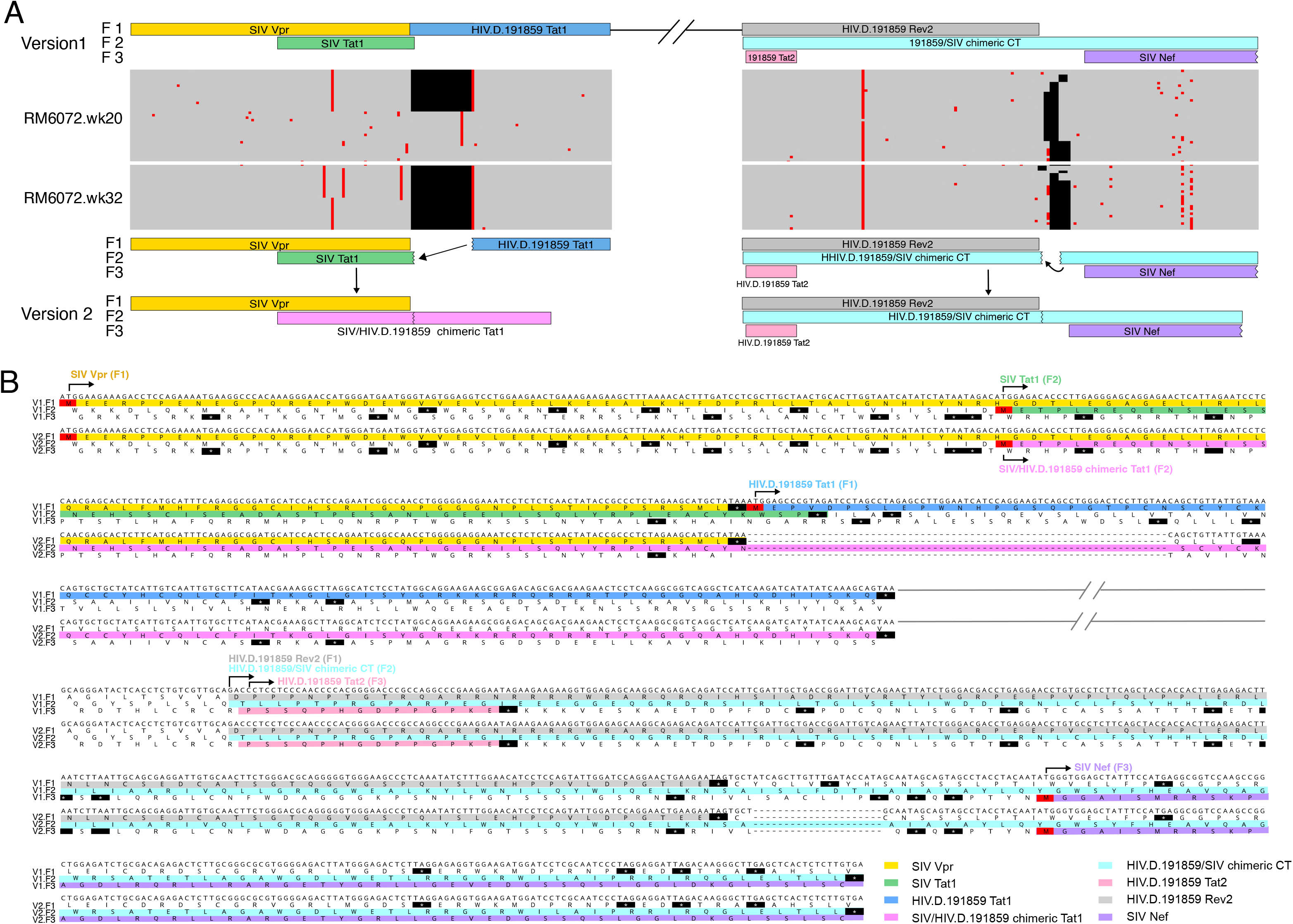
Spontaneous deletion of redundant HIV-1 and SIV *tat1 and env* gp41 sequences from the version 1 backbone in SHIV.C.CH505 infected monkeys. (**A**) Expanded segments of the *vpr – tat1* gene overlap and *tat2 – rev2 – env – nef gene* overlap from Fig. 1A are illustrated above Pixel plots (https://www.hiv.lanl.gov/content/sequence/pixel/pixel.html) of 39 single genome sequences from RM6072 20 weeks post-SHIV infection and 26 sequences 32 weeks post-SHIV infection. A 68 bp deletion of redundant sequences in *tat1* and a 21 bp deletion of redundant sequences in *env* gp41 rapidly becomes fixed in the evolving virus quasispecies. The version 2 backbone vector eliminates these redundant sequences. (**B**) Nucleotide and inferred amino acid sequences of the junctional regions of HIV-1 and SIV *tat1 and env* gp41 version 1 and 2 backbone vectors are shown, highlighting the differences in their designs.

**Figure S2.**
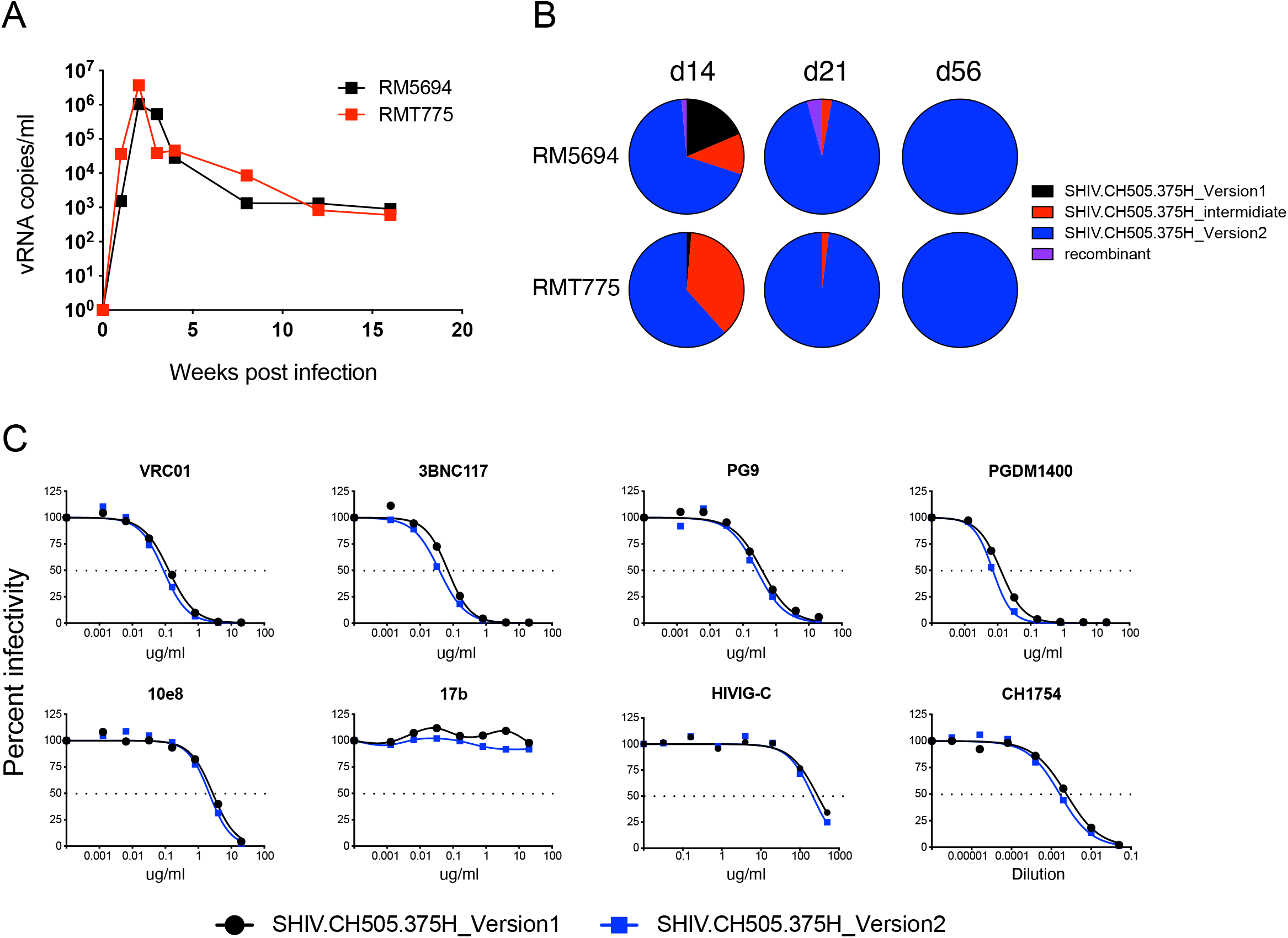
Comparison of replication efficiency, antigenicity and neutralization sensitivity of SHIV.C.CH505.375H derived from first (version 1) and second (version 2) generation construction schemes. Equal proportions of SHIV.CH505.375H versions 1, version 2 and an “intermediate version” were inoculated intravenously into two RMs and replication kinetics (**A**) and relative proportions of variants (**B**) were determined by plasma vRNA quantification and deep sequencing >1,000 viral genomes. Neutralization of version 1 and 2 SHIVs by prototypic human bNAb mAbs, the CD4-induced bridging sheet targeted mAb 17b, and polyclonal anti-HIV-1 IgG (HIVIG-C) and serum (CH1754) is depicted (**C**).

**Figure S3.**
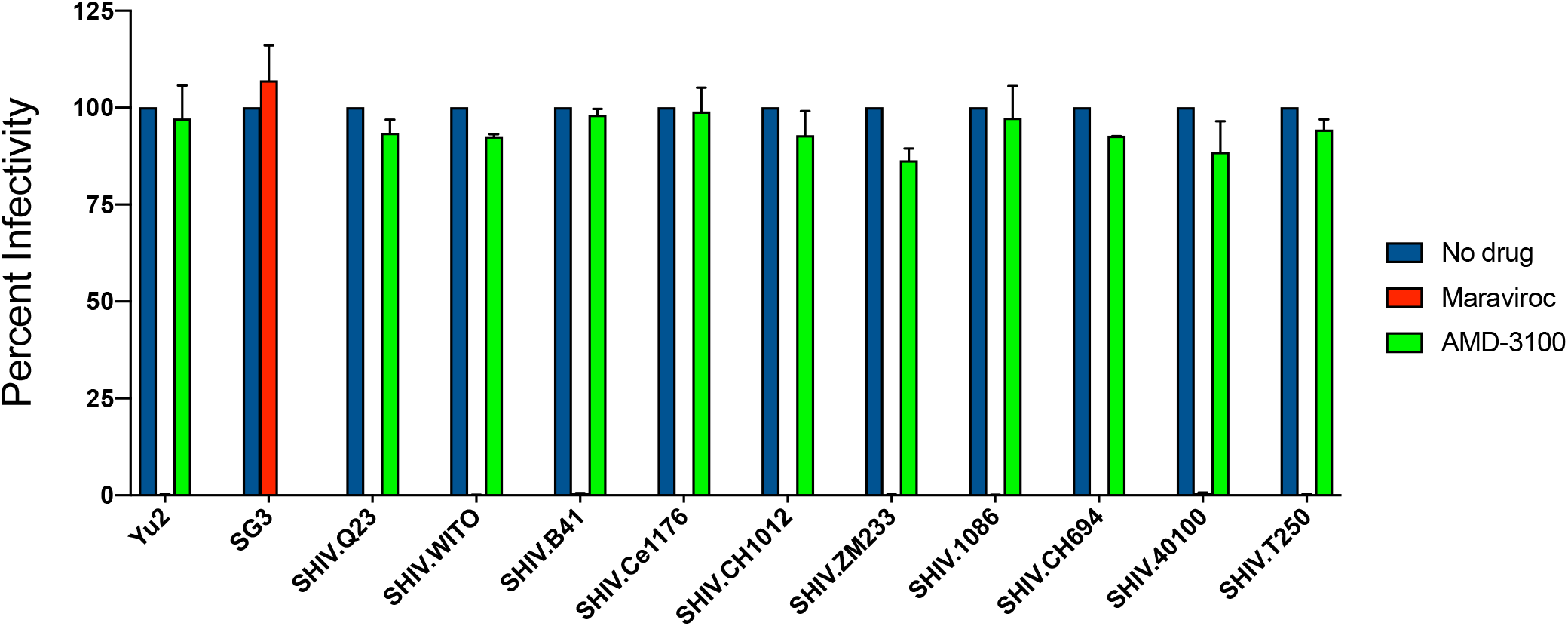
Inhibition of SHIV entry into TZMbl cells by the selective CCR5 and CXCR4 coreceptor inhibitors Maraviroc and AMD-3100, respectively. HIV-1 YU2 Env utilizes CCR5 exclusively for cell entry and HIV-1 SG3 Env utilizes CXCR4 exclusively. Maraviroc and AMD-3100 selectively abrogated cell entry of these viruses, as expected. Entry of the ten SHIVs bearing wildtype or rhesus-preferred Env375 alleles was inhibited by >99% by Maraviroc but minimally or not at all by AMD-3100, indicating a dependence on CCR5 for cell entry.

**Table S1.** Characteristics of SHIV inocula and clinical outcomes of 41 rhesus macaques

| SHIV strain | Sub-type | Wildtype 375 residue | Animal ID | 375 variant | Number of variants | Stock derivation | Dosage P27 ng* | Route<sup>Δ</sup> | CD8 Depletion | Peak VL<sup>§</sup> | Setpoint VL | Preferred 375 residues<sup>#</sup> | Clinical AIDS |
| --- | --- | --- | --- | --- | --- | --- | --- | --- | --- | --- | --- | --- | --- |
| CE1176 | C | 375-Ser | 6445 | S, M, H, Y, F, W | 6 | 293T | 300 | IV | No | 673,106 | 1,016 | H, W | No |
|  |  |  | 6448 | S, M, H, Y, F, W | 6 | 293T | 300 | IV | No | 482,934 | <250 | W, H | No |
|  |  |  | 6562 | S, M, H, Y, F, W | 6 | 293T | 300 | IV | No | 29,252 | <250 | W, F | No |
|  |  |  | 6563 | S, M, H, Y, F, W | 6 | 293T | 300 | IV | No | 894,432 | 20,608 | F, W | No |
| CH1012 | C | 375-Ser | T679 | S, M, H, Y, F, W | 6 | 293T | 300 | IV | No | 4,814,098 | <250 | Y, W, H | No |
|  |  |  | T680 | S, M, H, Y, F, W | 6 | 293T | 300 | IV | No | 4,759,788 | <250 | Y, W | No |
|  |  |  | 6928 | Y | 6 | 293T | 50 | IV | Yes | 42,350,830 | 8,830 |  | No |
|  |  |  | 6929 | Y | 6 | 293T | 50 | IV | Yes | 25,654,060 | 39,992 |  | Yes |
|  |  |  | 6930 | Y | 6 | 293T | 50 | IV | Yes | 14,495,480 | 10,656 |  | Yes |
| T250-4 | AG | 375-Ser | 6709 | S, M, H, Y, F, W | 6 | 293T | 300 | IV | No | 471,828 | <250 | H, W | No |
|  |  |  | 6716 | S, M, H, Y, F, W | 6 | 293T | 300 | IV | No | 1,744,646 | 5,708 | W, Y, F | No |
|  |  |  | 40701 | S, M, H, Y, F, W | 6 | 293T | 300 | IV | No | 932,684 | 0 | W, H | No |
|  |  |  | 6925 | Y, H, W, F | 1 | 293T | 200 | IV | Yes | 67,161,230 | 8,900 | H | No |
|  |  |  | 6926 | Y, H, W, F | 1 | 293T | 200 | IV | Yes | 9,600,904 | 29,196 | H | Yes |
|  |  |  | 6927 | Y, H, W, F | 1 | 293T | 200 | IV | Yes | 42,558,680 | 23,966 | H | No |
| Q23.17 | A | 375-Ser | 41298 | S, M, H, Y, F, W | 6 | 293T | 300 | IV | No | 122,560 | <250 | H, Y, F, W | No |
|  |  |  | 41852 | S, M, H, Y, F, W | 6 | 293T | 300 | IV | No | 173,420 | 7,680 | H, Y, F, W | No |
|  |  |  | T280 | H | 1 | 293T | 50 | IV | Yes | 29,118,930 | 29,000,000 |  | Yes |
| WITO4160 | B | 375-Thr | 40651 | T, S, M, H, Y, F, W | 7 | 293T | 350 | IV | No | 1,222,790 | 1,922 | W, Y | No |
|  |  |  | 40723 | T, S, M, H, Y, F, W | 7 | 293T | 350 | IV | No | 2,226,030 | 8,564 | W, Y | No |
|  |  |  | 40863 | T, S, M, H, Y, F, W | 7 | 293T | 350 | IV | No | 3,044,620 | 1,648 | W, Y | No |
| ZM233 | C | 375-Ser | 41412 | S, M, H, Y, F, W | 6 | 293T | 300 | IV | No | 51,052 | <250 | F, H, Y | No |
|  |  |  | 41537 | S, M, H, Y, F, W | 6 | 293T | 300 | IV | No | 408,840 | 2,390 | Y, H, F | No |
|  |  |  | 41712 | S, M, H, Y, F, W | 6 | 293T | 300 | IV | No | 364,598 | 7,068 | Y, F, H, M | No |
|  |  |  | 40717 | Y | 1 | 293T | 50 | IV | Yes | 1,432,284 | 18,936 |  | No |
|  |  |  | 11D042 | Y | 1 | 293T | 50 | IV | Yes | 61,300,980 | 88 |  | No |
|  |  |  | K12 | Y | 1 | 293T | 50 | IV | Yes | 4,445,834 | 2,072 |  | No |
| 1086 | C | 375-Ser | T682 | S, M, H, Y, F, W | 6 | 293T | 300 | IV | No | 190,208 | 11,062 | W | Yes |
|  |  |  | T683 | S, M, H, Y, F, W | 6 | 293T | 300 | IV | No | 682,242 | 2,090 | W, Y | No |
|  |  |  | T684 | S, M, H, Y, F, W | 6 | 293T | 300 | IV | No | 3,144,292 | <250 | W | No |
|  |  |  | T929 | W | 1 | 293T | 50 | IV | Yes | 2,679,424 | 20,138 |  | No |
|  |  |  | T930 | W | 1 | 293T | 50 | IV | Yes | 2,648,942 | 7,944 |  | No |
| CH0694 | C | 375-Thr | T931 | T, S, M, H, Y, F, W | 7 | 293T | 350 | IV | Yes | 7,778,446 | 106,716 | Y | No |
|  |  |  | T932 | T, S, M, H, Y, F, W | 7 | 293T | 350 | IV | Yes | 472,192 | 2,928 | Y, F | No |
|  |  |  | T933 | T, S, M, H, Y, F, W | 7 | 293T | 350 | IV | Yes | 3,718,208 | 119,506 | Y | No |
| B41 | B | 375-Ser | 41949 | S, M, H, Y, F, W | 6 | 293T | 300 | IV | No | 11,980,270 | 285,856 | H, W, Y | No |
|  |  |  | 42534 | S, M, H, Y, F, W | 6 | 293T | 300 | IV | No | 16,044,860 | 24,314 | H, W | No |
|  |  |  | 42579 | S, M, H, Y, F, W | 6 | 293T | 300 | IV | No | 4,332,634 | 387,742 | H,W | Yes |
| RV217.40100 | AE | 375-His | 40973 | S, M, H, Y, F, W | 6 | 293T | 300 | IV | No | 370,894 | 0 | W | No |
|  |  |  | 41193 | S, M, H, Y, F, W | 6 | 293T | 300 | IV | No | 81,162 | 0 | W | No |
|  |  |  | 41216 | S, M, H, Y, F, W | 6 | 293T | 300 | IV | No | 1,555,258 | 3,380 | W | No |

\* Total inoculum dose of virus as indicated with 50 ng of each Env375 variant
<sup>Δ</sup>IV – intravenous bolus by slow push
<sup>§</sup> VL – virus load (vRNA molecules/milliliter of plasma)
<sup>#</sup> Preferred Env375 allelic variants in plasma (H-His, W-Trp, F-Phe, Y-Tyr, M-Met)

